# Dynamic UFMylation governs cellular fitness by coordinating multi-organelle proteostasis

**DOI:** 10.64898/2026.03.27.714830

**Authors:** Guy B. Kunzmann, William E. Leiter, Sophie E. Durn, Amy M. Weeks, Jason R. Cantor

**Affiliations:** Morgridge Institute for Research, Madison, WI 53715, USA; Department of Biochemistry, University of Wisconsin-Madison, Madison, WI 53706, USA; Department of Chemistry, University of Wisconsin-Madison, Madison, WI 53706, USA; Carbone Cancer Center, University of Wisconsin-Madison, Madison, WI 53705, USA

## Abstract

Ubiquitin-fold modifier 1 (UFM1) is a ubiquitin-like protein (UBL) covalently attached to substrates through a dedicated enzymatic cascade (UFMylation) and removed by specific proteases. Despite a key role in endoplasmic reticulum (ER)-ribosome homeostasis, the basis by which this UBL supports cell fitness remains elusive, as the essentiality of UFMylation machinery varies widely across hundreds of cancer lines. Here, we trace a conditional dependence on the UFMylation pathway to the availability of alanine, an amino acid provided by human plasma-like medium but absent from most conventional synthetic media. We show that by facilitating the clearance of stalled ribosomes at the ER, dynamic UFMylation maintains cellular levels of glutamic-pyruvic transaminase 2 (GPT2), the primary enzyme responsible for de novo alanine synthesis in most human cancer lines. This buffering preserves the alanine pools required to sustain protein synthesis under alanine-restricted conditions. Beyond GPT2, *UFM1* deficiency leads to widespread proteomic remodeling that spans diverse processes, including mitochondrial translation. Our results reveal that despite primarily targeting ER-localized ribosomes, the UFMylation system orchestrates a multi-organelle proteostasis network whose client composition and contributions to cell fitness are shaped by intrinsic factors and nutrient conditions.

---

Genetic determinants of cell growth are shaped by an interplay between intrinsic and extrinsic factors^1^. Defining these determinants can help elucidate essential molecular functions and uncover potential therapeutic targets^2–5^. CRISPR-based screens across hundreds of cell lines have enabled the identification of core fitness genes required for cell growth in most cases, while also revealing that other genetic dependencies vary with the intrinsic diversity of human cancers^6–8^. However, despite increasing evidence that genetic dependencies are influenced by environmental factors^9–12^, there has been limited investigation into how growth conditions affect gene essentiality in proliferating human cells. Moreover, most CRISPR screens are still performed in conventional culture media that poorly resemble nutrient conditions encountered by cells in the human body^13^.

We previously developed human plasma-like medium (HPLM), a synthetic culture medium that contains over 60 components at concentrations selected to resemble those in healthy human blood^14^. To prepare complete HPLM-based media, we supplement with 10% dialyzed FBS (HPLM^+dS^), which provides various biomolecules needed for cell growth while minimizing the incorporation of undefined polar metabolites. In prior work, we demonstrated that medium composition can profoundly affect the genetic dependencies of human cancer cells^12^. Indeed, CRISPR screens in HPLM^+dS^ versus traditional media uncovered hundreds of conditionally essential genes. Deciphering the mechanisms underlying such conditional CRISPR phenotypes can yield critical insights into how specific proteins support cell growth and how nutrient conditions influence these contributions^12,15,16^.

UFM1 is a UBL covalently attached to target proteins by a dedicated E1-E2-E3 enzymatic cascade in a process known as UFMylation^17–19^. Similar to ubiquitination, this process is dynamic and reversible, as substrate-conjugated UFM1 can be detached via de-UFMylase-catalyzed reactions^20–23^. Despite a growing substrate landscape that spans various biological processes^24^, the molecular basis for how UFMylation supports cell fitness remains largely unresolved. Our genome-scale CRISPR screens revealed that the core UFMylation machinery – UFM1, the respective E1-E2-E3 enzymes, and the primary de-UFMylase – displayed greater essentiality in conventional media versus HPLM^+dS^. Here, we trace this differential dependence to alanine availability. We find that disruption of the UFMylation-de-UFMylation cycle leads to depletion of GPT2, the primary enzyme responsible for de novo alanine synthesis in most human cancer lines. We show that by facilitating the clearance of stalled ribosomes at the ER, dynamic UFMylation maintains GPT2 abundance, thereby preserving the alanine pools required to sustain protein synthesis under alanine-restricted conditions. Our data further demonstrate that disruption of this system leads to widespread proteomic remodeling beyond GPT2, including the selective depletion of mitoribosomal proteins, and that these effects vary with cell-intrinsic factors and nutrient conditions. Together, our findings reveal that the UFMylation system functions as a ribosome ‘collision counter’ and proteostatic buffer, uncovering an unforeseen link between dynamic UFMylation and a multi-organelle proteostasis network that governs cell fitness under metabolic and translational stress.

## RESULTS

### Conditional dependence on the UFMylation system is linked to alanine availability

UFM1 is translated as a precursor that undergoes proteolytic processing mediated by either UFM1-specific peptidase 1 (UFSP1) or UFSP2, exposing the C-terminal Gly83 required for target conjugation^25^ (Fig. 1a). Mature UFM1 is covalently attached to targets through a cascade sequentially catalyzed by the UFM1-dedicated E1 (UBA5), E2 (UFC1), and E3 (UFL1). Substrate-bound UFM1 is then enzymatically removed by UFSP1/2, completing the UFMylation-de-UFMylation cycle. RNA-seq data indicate that each component of this cycle is uniformly expressed across nearly 1,700 human cancer lines, except for *UFSP1*, which was nonetheless detected in 70% of these lines (Fig. 1b)^26^.

**Fig. 1.**
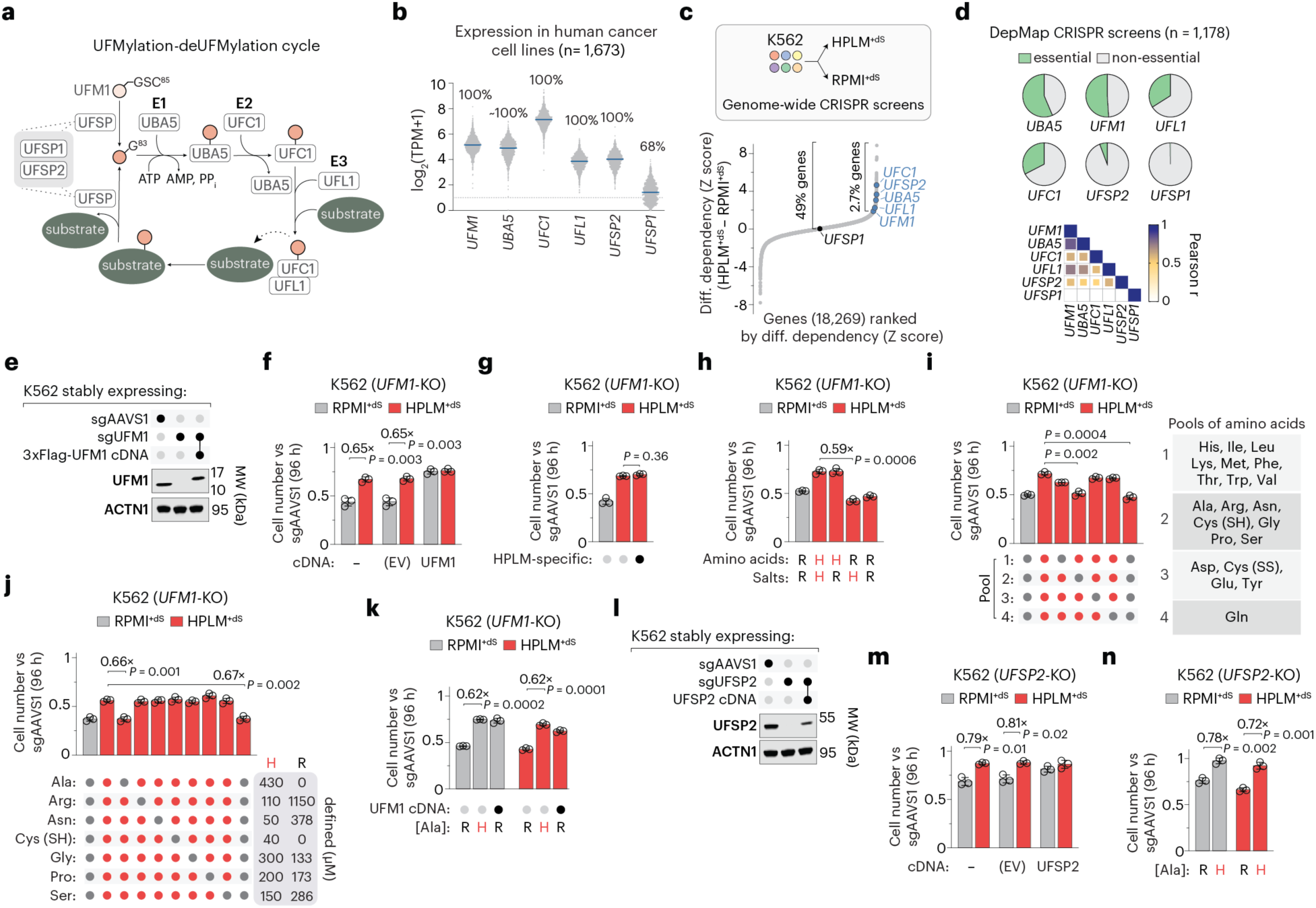
Conditional dependence on the UFMylation system is linked to alanine availability. (a) Schematic of the UFMylation-de-UFMylation cycle. UFM1 is processed by UFSP1/2, exposing the C-terminal Gly83 required for substrate conjugation. Mature UFM1 is covalently attached to substrates through an E1 (UBA5), E2 (UFC1), E3 (UFL1) enzymatic cascade. Substrate-bound UFM1 is detached by UFSP1/2-mediated de-UFMylation. (b) Distribution of mRNA transcript levels for core UFMylation machinery across human cancer lines from catalogued RNA-seq data^26^. Percentages indicate cell lines with log2(TPM + 1) > 1. Bolded lines denote the medians. (c) Genes ranked by differential dependency based on CRISPR screens in K562 cells^12^. Core UFMylation components are highlighted. (d) Dependency phenotypes for core UFMylation components in DepMap (top)^7^. Pearson correlation *(r)* matrix of co-dependency across cancer cell lines for the indicated gene pairs based on DepMap essentiality (probability of dependency > 0.5) (bottom). (e) Immunoblot for expression of UFM1 in *UFM1*-knockout and control cells. ACTN1 served as the loading control. (f-k) Relative growth of *UFM1*-knockout versus control cells (mean ± s.d., *n* = 3 biologically independent samples). Panel J (bottom right), defined levels of specified amino acids in HPLM and RPMI. EV, empty vector. (l) Immunoblot for expression of UFSP2 in *UFSP2*-knockout and control cells. ACTN1 served as the loading control. (m, n) Relative growth of *UFSP2*-knockout versus control cells (mean ± s.d., *n* = 3 biologically independent samples). In f-k, m, and n, Two-tailed Welch’s *t*-test comparing the respective mean ± s.d. (bar) versus mean ± s.d. (control cells) between bars. Values above brackets indicate fold change between bars. In h, i, and n, H, HPLM-defined concentration; R, RPMI-defined concentration. In i and j, red and dark gray dots denote HPLM- and RPMI-defined concentrations, respectively.

We previously performed genome-scale CRISPR screens in K562 chronic myeloid leukemia (CML) cells grown in either HPLM^+dS^ or RPMI prepared with HPLM-defined glucose and 10% dialyzed FBS (RPMI^+dS^)^12^. Analysis of these screens revealed that *UFM1*, *UBA5, UFC1, UFL1,* and *UFSP2* were among the top 3% of genes that displayed greater essentiality in RPMI^+dS^, whereas *UFSP1* was dispensable in both conditions (Fig. 1c). We first confirmed that each RPMI-essential component of the core UFMylation machinery was expressed in K562 cells and found that culture in HPLM^+dS^ versus RPMI^+dS^ did not affect the baseline abundance of these proteins (Extended Data Fig. 1a). Notably, while dependence on these components varies widely across cancer lines in DepMap^7^, those that scored as RPMI-essential exhibit strongly correlated dependency profiles (Pearson r = 0.48-0.82), highlighting the functional importance of dynamic UFMylation (Fig. 1d). K562 was not among the screened lines with a defined dependence on any of these genes, including *UFM1*, which is annotated as essential in roughly half of the nearly 1,200 cell lines in DepMap independent of lineage (Extended Data Fig. 1b, c).

To investigate the conditional CRISPR phenotype for UFMylation, we used a *UFM1*-targeting sgRNA to generate *UFM1*-knockout K562 clonal cells (Fig. 1e). Consistent with our screen results, *UFM1* knockout caused a nearly 40% stronger growth defect in RPMI^+dS^ versus HPLM^+dS^ relative to control cells transduced with an *AAVS1*-targeting sgRNA (Fig. 1f). Importantly, the expression of an sgRNA-resistant *UFM1* cDNA rescued this conditional defect, whereas transducing these cells with an equivalent empty vector did not.

We next sought to identify the difference in composition between HPLM and RPMI that may underlie conditional *UFM1* dependence. HPLM contains glucose, 10 salt ions, 10 vitamins (at RPMI-defined levels), 20 proteinogenic amino acids, and more than 30 additional metabolites absent from RPMI and most traditional synthetic media (Supplementary Table 1). To test whether any components within the latter set of HPLM-specific metabolites were responsible, we assessed the relative growth of *UFM1*-knockout cells in HPLM^+dS^ lacking this set but observed no impact compared to complete HPLM^+dS^ (Fig. 1g). We then evaluated relative growth in HPLM^+dS^ derivatives with the full sets of amino acids or salts adjusted to match RPMI-defined levels. Notably, culture in HPLM^+dS^ containing RPMI-defined amino acids impaired the growth of *UFM1*-knockout cells by an extent comparable to the defect in RPMI^+dS^ (Fig. 1h). By systematically adjusting the levels of amino acids in HPLM, first by subsets and then individually, we traced the conditional growth defect to alanine availability alone (Fig. 1i, j). Moreover, supplementing RPMI with HPLM-defined alanine (430 µM) rescued the conditional growth defect of *UFM1*-knockout cells (Fig. 1k). Alanine is the second most abundant amino acid in healthy human blood after glutamine but is absent from RPMI (Supplementary Table 1). Nonetheless, decreasing the alanine concentration in HPLM by up to 16-fold had negligible effects on the growth of *UFM1*-knockout cells, suggesting that even sub-physiologic alanine availability is sufficient to compensate for loss of the UFMylation system in these cells (Extended Data Fig. 1d).

While UFSP2 catalyzes de-UFMylation, our screens indicated that *UFSP2* shared a common conditional CRISPR phenotype with *UFM1* and the core E1-E2-E3 enzymes, consistent with the strong co-essentiality across this gene set in DepMap. Therefore, we investigated whether alanine availability similarly underlies conditional *UFSP2* dependence. Upon using a *UFSP2*-targeting sgRNA to generate clonal *UFSP2*-knockout K562 cells, we found that *UFSP2* deletion caused a 20% greater growth defect in RPMI^+dS^ versus HPLM^+dS^, which was rescued upon expressing an sgRNA-resistant *UFSP2* cDNA (Fig. 1l, m). We then assessed the relative growth of *UFSP2*-knockout cells in either HPLM^+dS^ lacking alanine or RPMI^+dS^ containing physiologic alanine (430 µM) and observed a similar impairment in each alanine-free condition (Fig. 1n). Collectively, these results show that alanine availability is necessary and sufficient to explain conditional dependence on dynamic UFMylation in K562 cells.

### The UFMylation system maintains GPT2 abundance

Since providing exogenous alanine restored growth in *UFM1*-knockout cells, we next sought to determine whether alanine levels in these cells were limiting relative to other amino acids. Cellular alanine pools were 5-fold smaller within control cells grown in RPMI^+dS^ versus HPLM^+dS^ (Extended Data Fig. 2a). *UFM1* deletion further reduced these levels by more than 10-fold specifically in RPMI^+dS^, while most other amino acids were only modestly depleted (30-50%) regardless of growth conditions (Fig. 2a). Despite not completely rescuing this RPMI-specific effect, expression of our *UFM1* cDNA increased alanine levels by 3-fold in the RPMI-cultured knockout cells. Together, *UFM1*-knockout cells grown in RPMI^+dS^ displayed a 70-fold reduction in alanine relative to those in HPLM^+dS^ (Fig. 2b). Similarly, loss of *UFSP2* reduced cellular alanine levels by 30-fold more in RPMI^+dS^ than in HPLM^+dS^, with only modest effects on other amino acid pools, an effect partially reversed by expression of our *UFSP2* cDNA (Extended Data Fig. 2b, c). We therefore reasoned that defects in the UFMylation pathway might disrupt de novo alanine production.

**Fig. 2.**
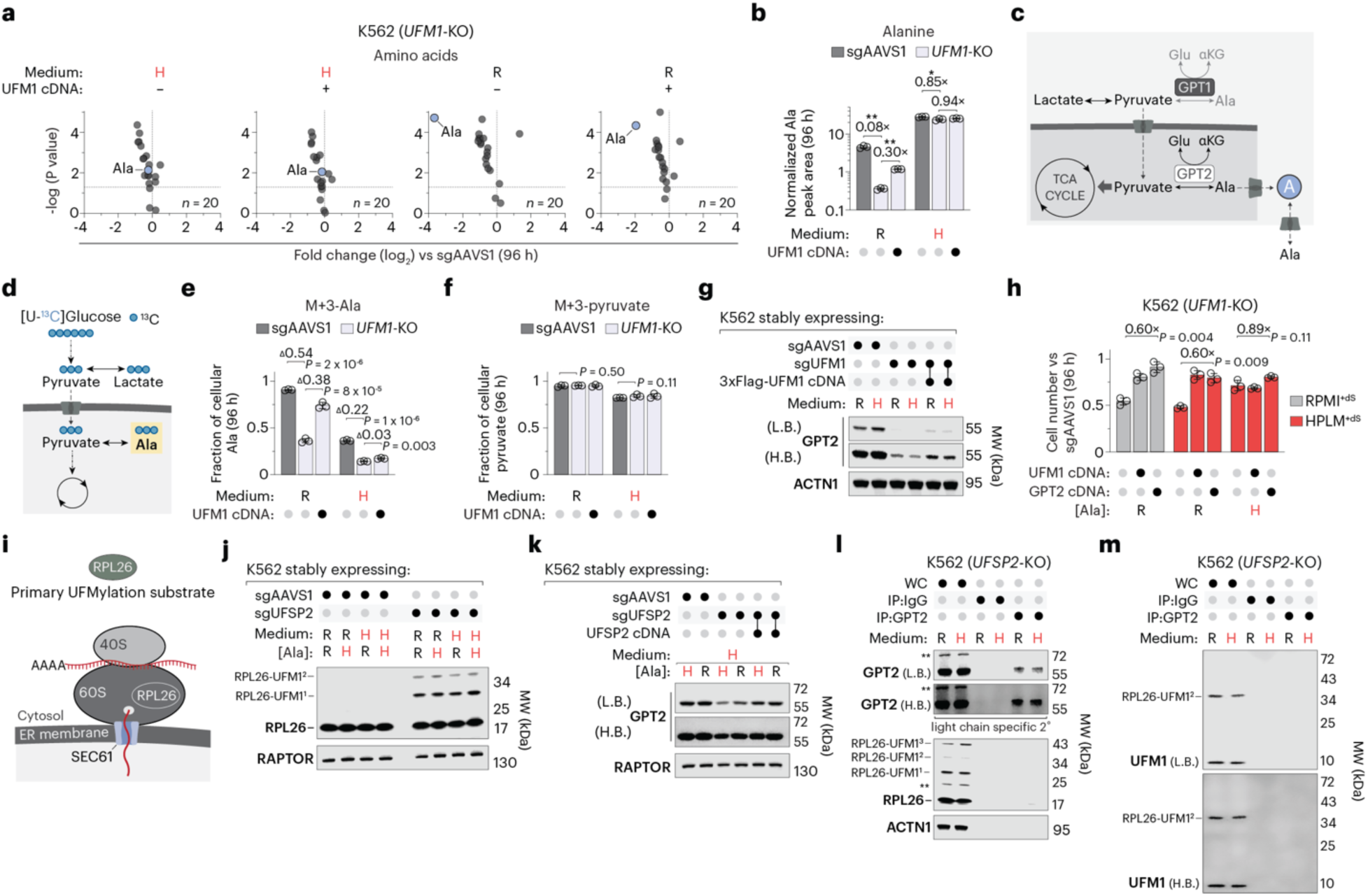
The UFMylation system maintains GPT2 abundance. (a) Relative amino acid levels in *UFM1*-knockout versus control cells (*n* = 3 biologically independent samples). Horizontal dotted lines mark a *P* value of 0.05 (y-axis). (b) Alanine levels in *UFM1*-knockout and control cells (mean ± s.d., *n* = 3 biologically independent samples). Two-tailed Welch’s *t*-test. \*\**P* < 0.005, \**P* < 0.01. (c) Schematic depicting exogenous alanine uptake and de novo synthesis from pyruvate. Most human cancer lines display near-selective expression of *GPT2* relative to *GPT1*^26^. (d) Schematic depicting the incorporation of ^13^C from [U-^13^C] glucose into alanine and lactate. (e, f) Fractional labeling of alanine (e) and pyruvate (f) in *UFM1*-knockout and control cells (mean ± s.d., *n* = 3 biologically independent samples). Two-tailed Welch’s *t*-test. Values above brackets indicate differences in fractional labeling between bars. (g) Immunoblot for expression of GPT2 in *UFM1*-knockout and control cells. ACTN1 served as the loading control. (h) Relative growth of *UFM1*-knockout versus control cells (mean ± s.d., *n* = 3 biologically independent samples). Two-tailed Welch’s *t*-test comparing the respective mean ± s.d. (bar) versus mean ± s.d. (control cells) between bars. (i) Schematic of RPL26, a component of the 60S subunit. RPL26 is the primary UFMylation substrate and is modified specifically on ER-localized ribosomes. (j, k) Immunoblots for expression of RPL26 (j) and GPT2 (k) in *UFSP2*-knockout and control cells. RAPTOR served as the loading control in both cases. (l, m) Immunoblots for expression of GPT2, RPL26 (l) and UFM1 (m) in whole-cell lysates (WC) and GPT2-immunopurified fractions (IP:GPT2) from *UFSP2*-knockout cells. IgG isotype control (IP:IgG) served as a negative control. ACTN1 served as the WC loading control. To prevent interference from the IgG heavy chain (∼50-55 kDa) with GPT2 signal (∼58 kDa), a light chain-specific secondary antibody was used for the GPT2 immunoblot. Asterisks in (l) denote non-specific bands. In b and h, values above brackets indicate fold change between bars. In h and j-m, H, HPLM-defined concentration; R, RPMI-defined concentration. In g and k-m, L.B., low brightness; H.B., high brightness.

Alanine synthesis in human cells is mediated by two GPT isoforms that can each catalyze the reversible conversion of pyruvate and glutamate to alanine and α-ketoglutarate, but differ in subcellular localization (GPT1, cytosol; GPT2, mitochondria) (Fig. 2c). Most human cancer cell lines, including K562, selectively express *GPT2* (Extended Data Fig. 2d)^26^. To determine whether *UFM1* deletion affects alanine synthesis, we assessed the incorporation of ^13^C into alanine in cells grown with [U-^13^C]-glucose (Fig. 2d). *UFM1* deletion reduced the fraction of alanine labeled with three ^13^C (M+3) by more than 50% in RPMI^+dS^ and by roughly 20% in HPLM^+dS^ (Fig. 2e). Consistent with the total pool effects, enforced expression of UFM1 partially rescued this defect in RPMI^+dS^. Notably, *UFM1* knockout did not affect the M+3-labeling of pyruvate or lactate in either condition (Fig. 2f and Extended Data 2e). Similarly, *UFSP2* deletion led to a 2-fold greater reduction in M+3 alanine labeling in RPMI^+dS^ (22%) versus HPLM^+dS^ (11%), with minimal impact on pyruvate or lactate labeling (Extended Data Fig. 2f-h). These results indicate that dynamic UFMylation promotes the diversion of glucose-derived carbon into alanine synthesis.

We next investigated whether impaired alanine production in UFMylation-deficient cells was linked to reduced GPT levels. Consistent with RNA-seq data, we previously observed that endogenous GPT1 was undetectable in K562 cells^12^. Therefore, we asked how loss of *UFM1* or *UFSP2* affects GPT2 abundance. Immunoblotting revealed that GPT2 was markedly depleted in both knockout lines regardless of nutrient conditions, an effect partially reversed by re-expressing the respective cDNA (Fig. 2g and Extended Data Fig. 2i). By contrast, qPCR showed that loss of either gene had minimal impact on *GPT2* mRNA levels, whereas *GPT2* knockout or overexpression considerably altered both mRNA and protein levels (Extended Data Fig. 2j, k). These results indicate that the UFMylation system regulates GPT2 abundance at the post-transcriptional level. Indeed, expression of a *GPT2* cDNA rescued the relative growth defect of *UFM1*-knockout cells in both RPMI^+dS^ and alanine-free HPLM^+dS^ by an extent comparable with enforced UFM1 expression (Fig. 2h). To assess whether mitochondrial localization of GPT activity was required for this rescue, we transduced our *UFM1*-knockout cells with a cytosol-restricted *GPT2* cDNA lacking the mitochondrial targeting signal (MTS) (Extended Data Fig. 2l, m). Similar to wild-type GPT2, expressing this MTS-deficient variant rescued the conditional growth defect, indicating that the alanine limitation was not compartmentalized (Extended Data Fig. 2n).

While an expanding number of reported substrates suggests UFMylation can influence protein stability in some cases^27–34^, evidence indicates that ribosomal protein uL24 (RPL26) – a component of the 60S subunit – is the primary target of this pathway, with UFM1 conjugation occurring exclusively on ER-localized ribosomes (Fig. 2i)^35–37^. Since UFM1 attachment is dynamic and often transient, *UFSP2* deletion can be leveraged to enrich for UFMylated substrates^35,37–39^. As anticipated, although we detected only unmodified RPL26 in our control cells, *UFSP2*-knockout cells displayed a robust accumulation of mono- and poly-UFMylated RPL26 regardless of culture in HPLM^+dS^ or RPMI^+dS^ (Fig. 2j). By contrast, immunoblot analysis for GPT2 in lysates from this knockout line revealed no evidence of similarly modified species (Fig. 2k).

To enrich for potential low-abundance conjugates, we immunopurified endogenous GPT2 from *UFSP2*-knockout cells but observed no modified GPT2 species or UFM1 signal in the purified eluent, despite detecting free UFM1 and UFMylated RPL26 in the corresponding lysates (Fig. 2l, m). Next, we engineered two *UFM1* cDNAs to encode: a conjugation-dead variant lacking the three C-terminal residues (UFM1ΔC3) and a mature variant exposing the reactive Gly83 (UFM1ΔC2). We then evaluated immunoblot patterns in lysates and Flag-purified eluents from *UFSP2*-knockout cells that expressed one of these variants or a control cDNA *(RAP2A)*. As expected, UFMylated RPL26 species were present across all lysates but exclusively enriched in the anti-Flag eluent from UFM1ΔC2-expressing cells (Extended Data Fig. 2o). However, we observed only unmodified GPT2 in these lysates, with a comparable background signal across the Flag-purified fractions. Flag signal analysis confirmed the presence of UFMylated RPL26 and an additional major band identified as a UBA5-UFM1 conjugate based on parallel immunoblotting (Extended Data Fig. 2p). Collectively, these results suggest that dynamic UFMylation indirectly regulates GPT2 abundance, likely at the translational level.

### The UFMylation system acts as a ribosome collision counter at the ER

We next asked whether established roles of the UFMylation pathway could provide a basis for its conditional essentiality, specifically involving RPL26. During translation, incidental ribosome stalling can occur due to a variety of factors – such as stable mRNA structures, problematic nascent peptide sequences, and nutrient limitation^40–43^ – leading to ribosome collisions that trigger RPL26 UFMylation on ER-localized ribosomes (Fig. 3a)^36,37^. Evidence indicates that this modification is mediated by a complex consisting of the UFM1-specific E3 ligase (UFL1), CDK5RAP3, and the ER-tethered adaptor DDRGK1^44,45^. RPL26 UFMylation promotes the clearance of these stalls through either ER-associated ribosome quality control (ER-RQC), which leads to the proteasomal degradation of arrested nascent chains, or selective autophagy of ER fragments (ER-phagy)^36,37,39,46,47^. Finally, RPL26 is de-UFMylated by UFSP2, which translocates from the cytosol via direct interaction with the ER-anchored ODR4^35,48^, enabling the release and recycling of stalled 60S subunits from the ER membrane^49,50^. This dynamic RPL26 UFMylation cycle is critical for maintaining ER-ribosome homeostasis.

**Fig. 3.**
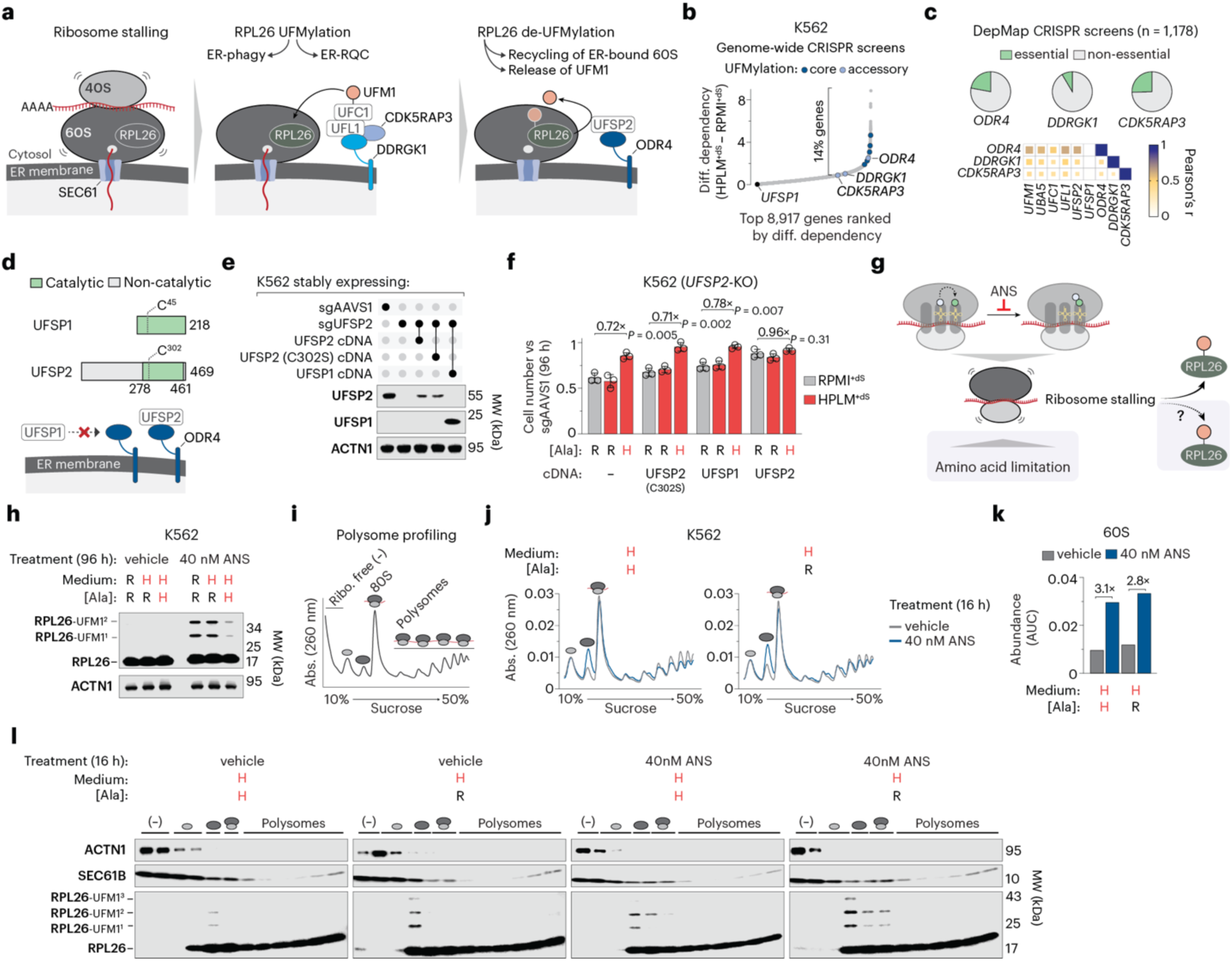
The UFMylation system acts as a ribosome collision counter at the ER. (a) Schematic of dynamic RPL26 UFMylation. Ribosome stalling (left) triggers RPL26-UFMylation on ER-localized ribosomes (middle). RPL26-UFM1 conjugation is mediated by a trimeric complex (UFL1, CDK5RAP3, and ER-tethered DDRGK1), leading to either ER-phagy or ER-RQC. RPL26 is then de-UFMylated by UFSP2, which is recruited to the ER membrane via interaction with ER-anchored ODR4, enabling the release and recycling of stalled 60S subunits from the ER membrane (right). (b) Subset of genes ranked by differential dependency based on CRISPR screens in K562 cells^12^. Accessory components of the ER-resident UFMylation machinery are highlighted. (c) Dependency phenotypes for (de)UFMylation accessory components in DepMap (top)^7^. Pearson correlation *(r)* matrix of co-dependency across cancer cell lines for the indicated gene pairs based on DepMap essentiality (probability of dependency > 0.5) (bottom). (d) Schematic of domain architecture for UFSP1 (top) and UFSP2 (bottom). UFSP2 contains a unique N-terminal domain that mediates recruitment to the ER membrane via interaction with ODR4. (e) Immunoblots for expression of UFSP1 and UFSP2 in *UFSP2*-knockout and control cells. ACTN1 served as the loading control. (f) Relative growth of *UFSP2*-knockout versus control cells (mean ± s.d., *n* = 3 biologically independent samples). Two-tailed Welch’s *t*-test comparing the respective mean ± s.d. (bar) versus mean ± s.d. (control cells) between bars. (g) Schematic of ribosome stalling drivers. Anisomycin (ANS) is a global elongation inhibitor that binds the 60S subunit and triggers RPL26 UFMylation. Amino acid restriction induces codon-specific stalls via depletion of charged tRNAs. (h) Immunoblot for expression of RPL26 in ANS- and vehicle-treated cells. ACTN1 served as the loading control. (i) Schematic of a polysome profile depicting the distribution of free ribosomal subunits (40S, 60S), monosomes (80S), and polysomes across a sucrose gradient. Ribo., ribosome. (j) Polysome profiles from ANS- and vehicle-treated cells. (k) Abundance of free 60S in ANS- and vehicle-treated cells, quantified by area under the curve (AUC) analysis of polysome profiles. (l) Immunoblots for expression of SEC61B and RPL26 in sucrose gradient fractions collected from ANS- and vehicle-treated cells. In f, h, and j-l, H, HPLM-defined concentration; R, RPMI-defined concentration. In f and k, values above brackets indicate fold change between bars.

To examine whether ER localization contributes to conditional *UFM1* dependence, we first assessed the composition of the active UFMylation machinery in HPLM^+dS^ and RPMI^+dS^. Using *UFM1*-knockout cells expressing either wild-type UFM1, the conjugation-dead variant (ΔC3), or a control bait (RAP2A), we performed Flag-based affinity purification-mass spectrometry (AP-MS) (Extended Data Fig. 3a). Notably, the only prey proteins significantly enriched (FDR < 0.05, fold-change > 10) in wild-type UFM1 samples relative to the control bait were UBA5, UFC1, UFL1, DDRGK1, and CDK5RAP3, regardless of culture conditions (Extended Data Fig. 3b and Supplementary Table 3). By contrast, UBA5 was the only such hit captured by UFM1ΔC3 in both conditions. These data are consistent with evidence that UFM1 and UBA5 can interact non-covalently and further support the notion that covalent conjugation is required for stable association with the E2 enzyme and the trimeric E3 complex^49–52^. Together, these results suggest that the UFMylation machinery is assembled and poised at the ER independent of nutrient conditions.

Next, we asked whether our CRISPR screen data supported the possibility that ER localization is critical to the conditional essentiality of the UFMylation system. Similar to the core machinery, *ODR4* scored as a strong RPMI-essential hit, while *DDRGK1* and *CDK5RAP3* ranked among the top 14% of genes with greater essentiality in RPMI^+dS^ versus HPLM^+dS^ (Fig. 3b)^12^. While dependency profiles for these three accessory components vary across DepMap, *ODR4* displays stronger co-essentiality with the core components (mean Pearson r = 0.59) than either *DDRGK1* (0.43) or *CDK5RAP3* (0.37) (Fig. 3c). These data suggest that assembly of the UFSP2-ODR4 complex for ER-localized UFM1 removal is particularly crucial to the role of this system in supporting cell growth.

While UFSP1 and UFSP2 can both mediate UFM1 maturation and substrate de-UFMylation, UFSP2 contains a unique N-terminal domain that enables recruitment to the ER membrane via ODR4 (Fig. 3d)^20,22,23,48,53^. Therefore, we asked whether conditional *UFSP2* dependence is linked to the ER localization of UFSP activity. To test this possibility, we engineered a protease-dead *UFSP2* cDNA by mutating the catalytic cysteine residue (C302S)^20,21,54^. Enforced expression of either this catalytically inactive UFSP2 or wild-type UFSP1 failed to rescue the conditional growth defect of *UFSP2*-knockout cells, suggesting that ER-resident de-UFMylase activity is necessary to sustain cell growth in RPMI^+dS^ or alanine-free HPLM^+dS^ (Fig. 3e, f).

Amino acid restriction can deplete charged tRNAs, inducing codon-specific ribosome stalls, whereas the translation inhibitor anisomycin drives a global elongation block^40,55–57^. In both cases, ribosome stalling can lead to ribosome collisions. While prior work established that anisomycin triggers RPL26 UFMylation, whether individual amino acid restriction also promotes this modification remains unexplored (Fig. 3g)^36,37^. Since our data indicated that ER localization is critical for how the UFMylation system supports cell growth, we reasoned that increased dependence on this system under alanine-restricted conditions may be linked to its known role in ER-ribosome homeostasis. To test this, we first assessed RPL26 UFMylation levels in K562 cells treated with anisomycin (40 nM). As anticipated, RPL26-UFM1 conjugates were detected upon immunoblot analysis of whole-cell lysates only in anisomycin-treated cells, with markedly higher accumulation in RPMI^+dS^ and alanine-free HPLM^+dS^ compared to complete HPLM^+dS^ (Fig. 3h). Given that such whole-cell snapshots lack the resolution to distinguish specific ribosomal states, we used polysome profiling to determine the distribution of modified RPL26 (Fig. 3i). Consistent with previous work, anisomycin treatment caused a buildup of free 60S subunits^58,59^, a relative impact unaffected by alanine restriction (Fig. 3j, k). Notably, while baseline RPL26 modification was undetectable in whole-cell lysates, polysome analysis revealed that UFMylated RPL26 co-sedimented with 60S subunits in vehicle-treated cells, reaching considerably higher levels in alanine-free HPLM^+dS^ (Fig. 3l). Together, while anisomycin was a more potent stimulus than alanine restriction alone, the combination of both stressors led to the highest enrichment of UFMylated RPL26, indicating the ER-resident UFMylation system effectively functions as a ‘collision counter’ that responds proportionally to the total ribosome collision burden. These results demonstrate that alanine restriction can trigger ribosome UFMylation and suggest that alanine limitation drives an increased dependence on this system to resolve stalled ribosomes at the ER.

### Dynamic UFMylation is conditionally required to maintain protein synthesis rates

Guided by our data establishing that the UFMylation system senses stress-induced ribosome collisions, we next asked how impaired (de)UFMylation affects the distribution of ribosomal subunits, a proxy for unresolved stalls, upon alanine restriction (Fig. 4a, b). Polysome profiling revealed that loss of *UFM1* or *UFSP2* alone had little impact on free 40S levels in complete HPLM^+dS^, whereas alanine restriction led to a 10% reduction in control cells (Fig. 4c). Deleting either gene exacerbated this effect, with 40S levels reduced by 20-30% relative to control cells in alanine-replete HPLM^+dS^. By contrast, while alanine availability alone did not alter free 60S subunit levels, loss of *UFM1* or *UFSP2* drove a 50-70% increase in free 60S – an effect amplified under alanine restriction, resulting in 60S levels 2- to 3-fold higher than those for control cells in complete HPLM^+dS^ (Fig. 4d). We also found that 80S monosome levels were unaffected by either genetic deletion but decreased by 20-30% upon alanine removal (Fig. 4e). Together, alanine restriction increased the 40S:80S ratio by 30% in control cells, whereas loss of *UFM1* or *UFSP2* had little impact on this ratio, regardless of alanine availability, relative to the baseline in complete HPLM^+dS^ (Extended Data Fig. 4a). However, while either genetic deletion or alanine removal independently elevated the 60S:80S ratio by 50-70%, their combination led to a 3-fold increase across both knockout lines relative to control cells in alanine-replete HPLM^+dS^, mirroring the additive accumulation of RPL26 UFMylation in response to anisomycin under alanine restriction (Fig. 4f). Overall, these results demonstrate that impaired (de)UFMylation causes a 60S-specific quality control defect, consistent with the reported role of this system in recycling ER-localized 60S subunits^49,50^. By contrast, alanine limitation distinctly induces a general translational block, with these dual stressors leading to a synergistic collapse of ribosome homeostasis (Fig. 4g).

**Fig. 4.**
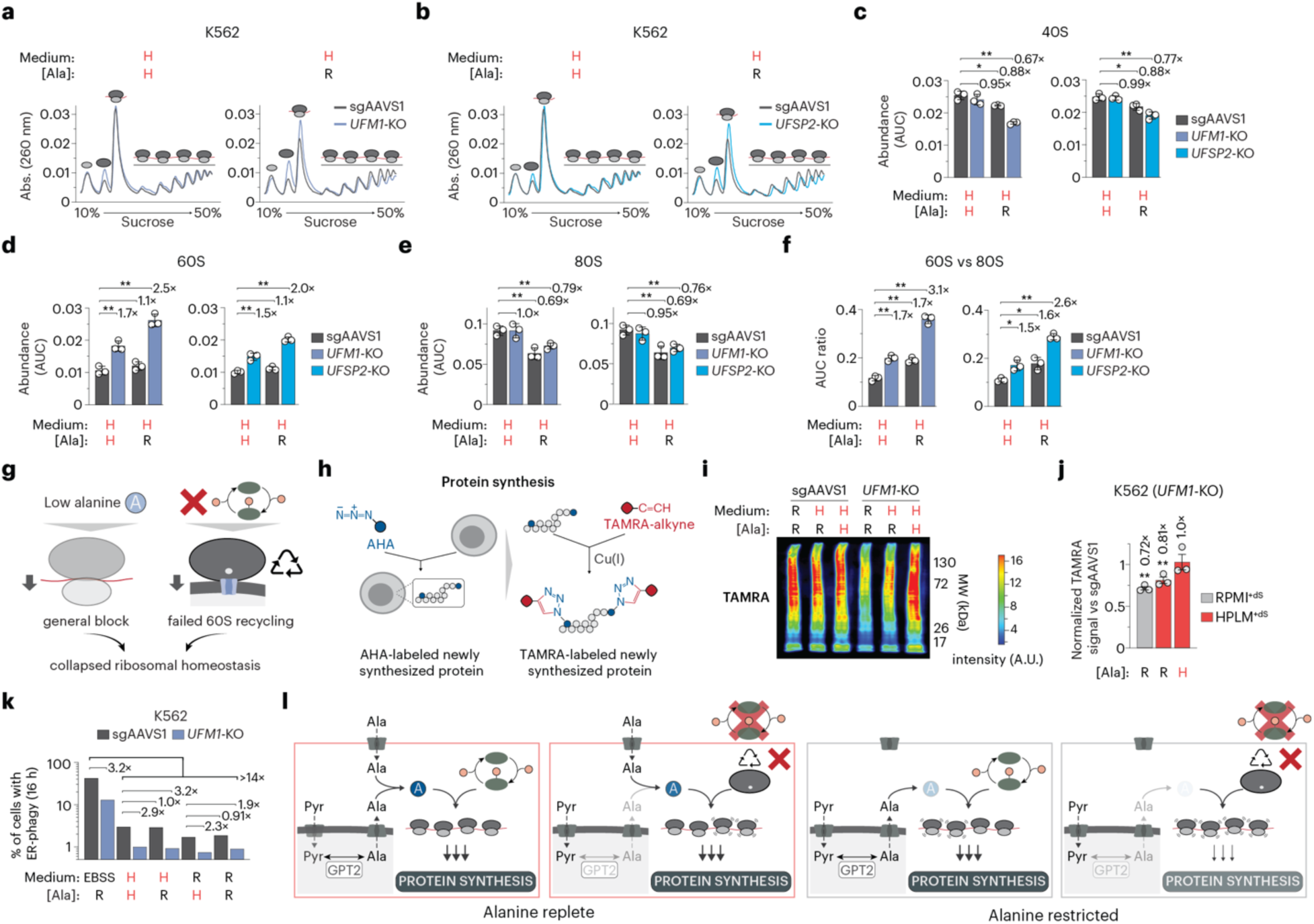
Dynamic UFMylation is conditionally required to maintain protein synthesis rates. (a, b) Polysome profiles from *UFM1*-knockout (a), *UFSP2*-knockout (b), and control cells. (c-e) Abundances of free 40S (c), free 60S (d), and 80S monosomes (e) in *UFM1*-knockout, *UFSP2*-knockout, and control cells, quantified by area under the curve (AUC) analysis of polysome profiles (mean ± s.d., *n* = 3 biologically independent samples). (f) Ratio of free 60S subunits to 80S monosomes in *UFM1*-knockout, *UFSP2*-knockout, and control cells (mean ± s.d., *n* = 3 biologically independent samples). (g) Proposed model of collapsed ribosomal homeostasis under additive translational stress. Loss of *UFM1* or *UFSP2* triggers a 60S-specific quality control defect (impaired recycling of ER-localized 60S subunits), while alanine limitation independently drives a general translational block. (h) Schematic depicting the evaluation of protein synthesis rates via metabolic pulse-labeling. Cells are pulsed with L-azidohomoalanine (AHA) to label nascent proteins (left), followed by conjugation to TAMRA-alkyne via copper (Cu)-catalyzed click chemistry (right). (i) Pseudocolor immunoblot of TAMRA signal in *UFM1*-knockout and control cells. Signal intensity is represented by the corresponding colorimetric scale. (j) Relative TAMRA signal in *UFM1*-knockout versus control cells (mean ± s.e.m., *n* = 3 biologically independent samples). For each replicate, raw intensities were normalized by total protein and then scaled to the maximum normalized value within that replicate. (k) Quantification of ER-phagy flux in *UFM1*-knockout and control cells. (l) Proposed model for conditional dependence on the UFMylation system based on relative alanine availability. In a-f, j, and k, H, HPLM-defined concentration; R, RPMI-defined concentration. In c-f and j, Two-tailed Welch’s *t*-test. \*\**P* < 0.01, \**P* < 0.05. In c-f and k, values above brackets indicate fold change between bars.

Given this global imbalance in ribosomal subunits, we reasoned that conditional dependence on the UFMylation system might stem from impaired translational output. While we observed modest reductions in the polysome fraction across both knockout lines specifically under alanine restriction (Fig. 4a, b), we sought to test this idea directly by quantifying protein synthesis rates. Using metabolic pulse-labeling, we cultured cells in methionine-free media containing L-azidohomoalanine (AHA) to label nascent proteins before conjugating the labeled proteins to TAMRA-alkyne via copper-catalyzed click chemistry (Fig. 4h)^60^. Immunoblot analysis revealed that loss of *UFM1* reduced protein synthesis rates (TAMRA signal normalized to total protein) by 20-30% relative to control cells in either RPMI^+dS^ or alanine-free HPLM^+dS^ (Fig 4i, j and Extended Data Fig. 4b). In contrast, *UFM1* deletion had no impact on AHA incorporation in complete HPLM^+dS^.

Since UFMylation promotes the resolution of stalled ribosomes by mediating ER-RQC or ER-phagy, we next asked whether *UFM1* deletion differentially influences ER-phagy in response to alanine restriction. To examine this, we used the ER-autophagy tandem reporter (EATR) system to evaluate ER-phagy flux^61^. As expected, incubation in Earle’s buffered saline solution (EBSS) induced robust ER-phagy in nearly 40% of control cells, a response 3-fold higher than in *UFM1*-knockout cells (Fig. 4k). However, while *UFM1* deletion reduced ER-phagy by a similar extent in cells cultured in HPLM^+dS^ or RPMI^+dS^, these effects were independent of alanine availability. Moreover, total ER-phagy flux was negligible (< 3%) across all conditions relative to EBSS starvation, regardless of *UFM1* expression.

Collectively, our results suggest a model in which loss of (de)UFMylation leads to defects in ER-associated 60S recycling and a marked reduction in GPT2 abundance. However, these defects are exacerbated under alanine restriction, where cellular alanine levels fall below a critical threshold, driving a surge in ribosome collisions. In turn, the ER-RQC pathway is overwhelmed by this cumulative stalling burden, thereby limiting the pool of available ribosomal subunits and impairing global protein synthesis rates required to sustain cell growth (Fig. 4l).

### Broad amino acid restriction does not impose increased dependence on UFMylation

Among proteinogenic amino acids, only alanine is uniquely defined in HPLM relative to RPMI. Therefore, we investigated whether restricting other amino acids might similarly induce a conditional growth defect in *UFM1*-knockout cells, focusing on non-essential amino acids (NEAAs) (Fig. 5a). Because SLC7A11, which forms a complex with SLC3A2 to mediate cystine-glutamate exchange, is a reported UFMylation substrate^33^, we also included the semi-essential cyst(e)ine in our panel (Fig. 5b). We systematically generated HPLM derivatives lacking either cyst(e)ine or single NEAAs, except for serine and glycine, which were removed in tandem since they can be metabolically interconverted. As expected, restricting individual NEAAs did not impact the growth of control cells, while the combined removal of serine/glycine or cysteine/cystine reduced total growth by roughly one or two doublings, respectively (Fig. 5c). Notably, *UFM1*-knockout cells displayed a relative growth defect only in the alanine-free derivative (Fig. 5d). These results suggest that restricting other amino acids within our panel did not trigger ribosome stalling sufficient to exceed the capacity of the ER-RQC pathway in UFMylation-deficient cells. Nonetheless, the limiting thresholds required to induce such persistent stalling likely depend on codon usage and the relative cellular abundance of specific amino acids^40^.

**Fig. 5.**
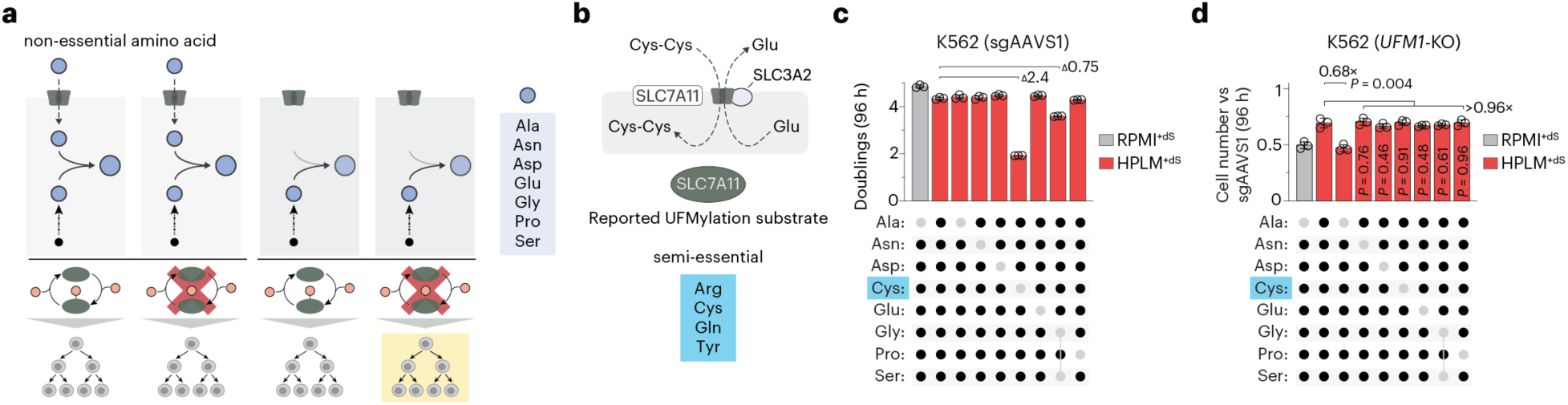
Broad amino acid restriction does not impose increased dependence on UFMylation. (a) Schematic depicting the integration of exogenous uptake and de novo synthesis into cellular pools of non-essential amino acids (NEAAs) (left, top). Experimental setup assessing the impact of specific NEAA restrictions on the relative growth of UFMylation-deficient cells (left, bottom). List of human NEAAs (right). (b) Schematic of system xc-, a heterodimeric complex of SLC7A11 and SLC3A2 that mediates the exchange of intracellular glutamate for extracellular cystine. SLC7A11 is a reported UFMylation substrate^33^. (c) Total population doublings for control cells (mean ± s.d., *n* = 3 biologically independent samples). Values above brackets indicate difference in total doublings. (d) Relative growth of *UFM1*-knockout versus control cells (mean ± s.d., *n* = 3 biologically independent samples). Two-tailed Welch’s *t*-test comparing the respective mean ± s.d. (bar) versus mean ± s.d. (control cells) between bars.

### UFM1 deletion remodels the global proteome

Although impaired (de)UFMylation depleted GPT2 levels regardless of growth in HPLM^+dS^ or RPMI^+dS^, we reasoned that this pathway might influence proteostasis beyond GPT2 regulation, with broad consequences that may also vary by nutrient conditions (Fig. 6a). To test this, we performed label-free untargeted proteomics to assess protein levels in *UFM1*-knockout versus control cells (Fig. 6b). First, we observed that culture in HPLM^+dS^ relative to RPMI^+dS^ had a negligible impact on the baseline proteome, with only five of nearly 5,000 quantified proteins significantly altered (FDR < 0.05, fold-change > 2) (Fig. 6c). By contrast, *UFM1* deletion induced widespread effects, though the fraction of differentially regulated proteins was roughly 2-fold larger in RPMI^+dS^ (4.8% down, 4.2% up) than in HPLM^+dS^ (2.0% down, 2.6% up) (Fig. 6d). However, GPT2 fell below the limit of detection across these datasets, perhaps reflecting its low relative expression levels.

**Fig. 6.**
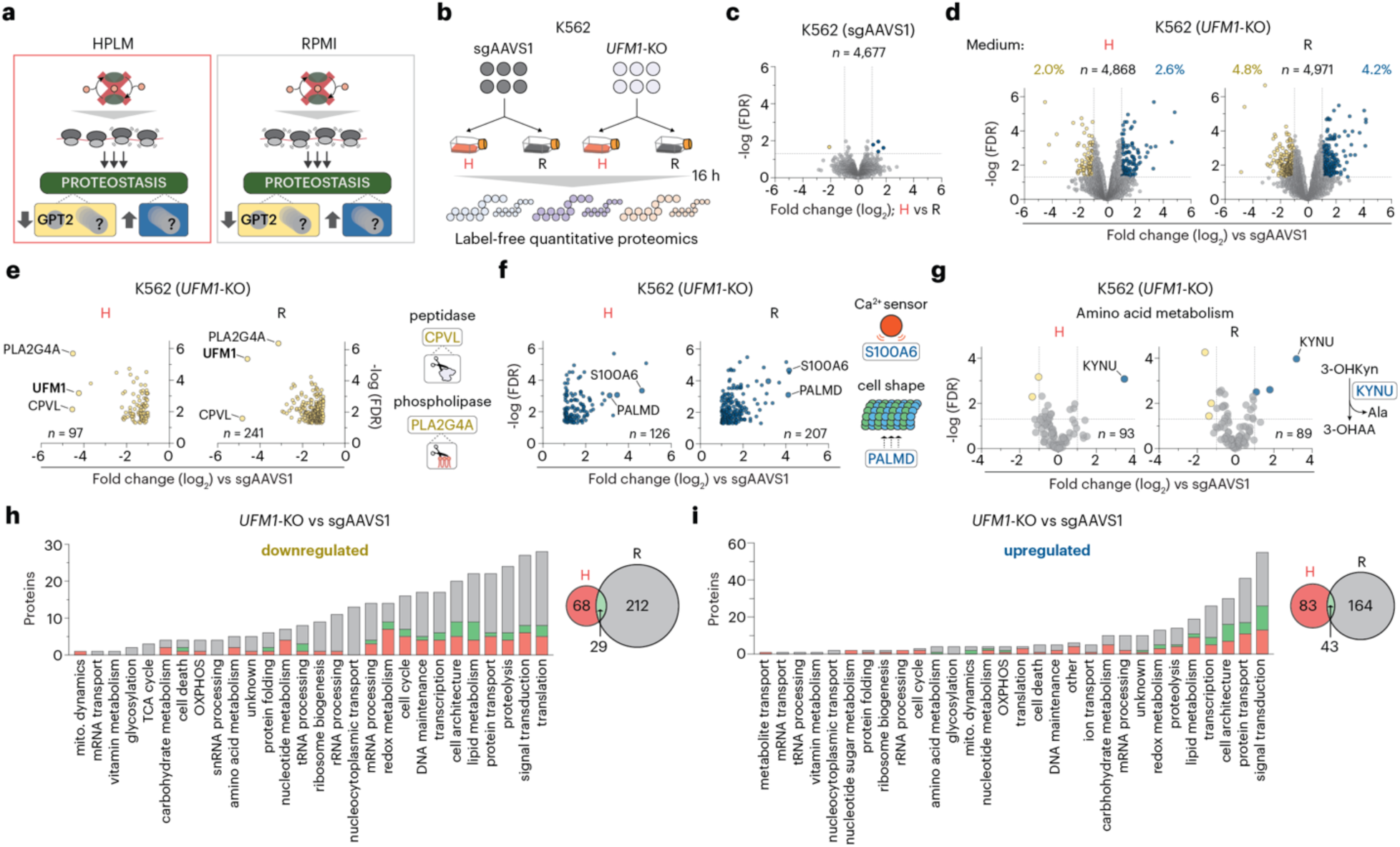
*UFM1* deletion remodels the global proteome. (a) Schematic depicting the hypothesized impact of impaired UFMylation on proteostasis beyond GPT2 depletion. (b) Workflow for quantitative label-free proteomics in *UFM1*-knockout and control cells. (c) Relative protein levels for control cells in HPLM^+dS^ versus RPMI^+dS^ (*n* = 6 biologically independent samples). (d-g) Relative protein levels in *UFM1*-knockout versus control cells (*n* = 6 biologically independent samples). (h, i) Curated biological processes for total downregulated (h) and upregulated (i) protein hits in *UFM1*-knockout versus control cells (left). Venn diagrams showing the overlap of these hits between HPLM^+dS^ and RPMI^+dS^ (right). In c-g, log2 fold change and FDR values were determined using FragPipe-Analyst^85^. In c, d, and g, dotted lines denote a fold change of ± 2 (x-axis) and an FDR of 0.05 (y-axis). In d, percentages denote differentially abundant proteins among the total quantified proteome.

We next examined the top differentially abundant proteins in *UFM1*-knockout versus control cells, focusing on hits shared across conditions. Beyond UFM1 itself, the two most markedly depleted proteins in both cases were the putative secretory enzyme carboxypeptidase vitellogenic-like (CPVL)^62^ and phospholipase A2 group IVA (PLA2G4A), which catalyzes arachidonic acid release upon Ca^2+^-mediated translocation to subcellular membranes (Fig. 6e). Conversely, the Ca^2+^ sensor S100A6 and palmdelphin (PALMD) – a protein involved in cell shape control, which is regulated in part by Ca^2+^ levels^63^ – were among the top upregulated proteins in each case (Fig. 6f). While none of these hits have been formally characterized in the context of UFMylation, they are all functionally tied to ER homeostasis. Of note, large-scale interactome mapping (BioPlex) previously identified S100A6 as a candidate UFM1 interactor in HEK293T cells^64^.

Next, we asked whether *UFM1* deletion altered the levels of other proteins that participate in amino acid metabolism beyond GPT2. While a focused analysis of nearly 100 such proteins – including a subset of almost 20 enzymes involved in amino acid synthesis – revealed only a few largely modest and non-overlapping hits, kynureninase (KYNU) ranked among the top 3% of upregulated proteins in each condition (Fig. 6g and Extended Data Fig. 5a). KYNU catalyzes the conversion of 3-hydroxykynurenine to 3-hydroxyanthranilic acid (3-OHAA), releasing alanine as a byproduct. Independent of downstream 3-OHAA metabolism, the elevated KYNU levels in *UFM1*-knockout cells may represent a compensatory response to GPT2 depletion, serving as an alternative, albeit seemingly inefficient, source of alanine. Together, these data further suggest that the UFMylation system is not a broad regulator of amino acid metabolism; rather, its disruption confers a specific vulnerability to alanine restriction through a targeted impact on GPT2 abundance.

Given that (de)UFMylation supports cell growth by facilitating ER-RQC, we also performed a focused assessment of nearly 60 RQC-associated proteins. Notably, among the minimal changes observed across this panel, we found that nuclear export mediator factor (NEMF) – which in part helps coordinate ER-RQC with the UFMylation system^47^ – was depleted by roughly 2-fold in *UFM1*-knockout cells across both conditions, falling just short of our defined hit thresholds in HPLM^+dS^ (Extended Data Fig. 5b). This suggests that functional UFMylation exerts both direct and indirect influence on the integrity of the ER-RQC pathway.

Although a small subset of the most differentially regulated proteins converged regardless of culture in HPLM^+dS^ or RPMI^+dS^, we manually categorized the entire set of hits by biological process to more broadly assess the consequences of *UFM1* deletion (Supplementary Table 4). Our analysis revealed that these proteins span a diverse range of functions and, further, that the overall proteomic signatures showed minimal overlap, with just 9% of downregulated and 15% of upregulated hits shared across conditions (Fig. 6h, i). Translation was the most highly represented process among the combined set of downregulated hits, while beyond the general category of signal transduction, protein transport and cellular architecture were the most frequent among the upregulated set. Together, these results suggest that impaired UFMylation markedly remodels the proteome, with comprehensive effects that are further shaped by nutrient conditions.

### UFM1 deletion alters mitochondrial proteostasis

Given the broad range of proteomic changes, we used Gene Ontology (GO) analysis to identify significantly overrepresented biological processes and cellular components. Pooling the upregulated hits across both conditions, we identified intracellular protein transport as the only enriched process (FDR < 0.05) (Extended Data Fig. 6a). Overrepresented components were primarily associated with the secretory pathway or cellular architecture – including focal adhesions, the ER-Golgi intermediate compartment, transport vesicles, the oligosaccharyltransferase complex, and secretory granule membranes – with the latter two categories composed only of RPMI-specific hits. Consistent with these data, a focused analysis of nearly 20 proteins involved in co-translational targeting to the ER revealed that *UFM1* deletion increased the mean abundance across this panel by 40% (two hits) and 70% (nine hits) in HPLM^+dS^ and RPMI^+dS^, respectively (Extended Data Fig. 6b). These results indicate that the loss of functional UFMylation leads to an accumulation of proteins linked to ER homeostasis, with effects exacerbated by the elevated translation stress in RPMI^+dS^.

We next performed a similar GO enrichment analysis for the pooled set of downregulated hits. Unexpectedly, mitochondrial translation was the only enriched process, with an FDR several orders of magnitude lower than that observed for the protein transport category described above (Fig. 7a). The most highly enriched cellular components included the mitochondrial matrix, the inner membrane, and mitoribosomal subunits. This analysis also identified components critical for ribosome biogenesis (nuclear envelope, nuclear pore, and the t-UTP complex), which were enriched to a lesser extent and consisted almost entirely of RPMI-specific hits.

**Fig. 7.**
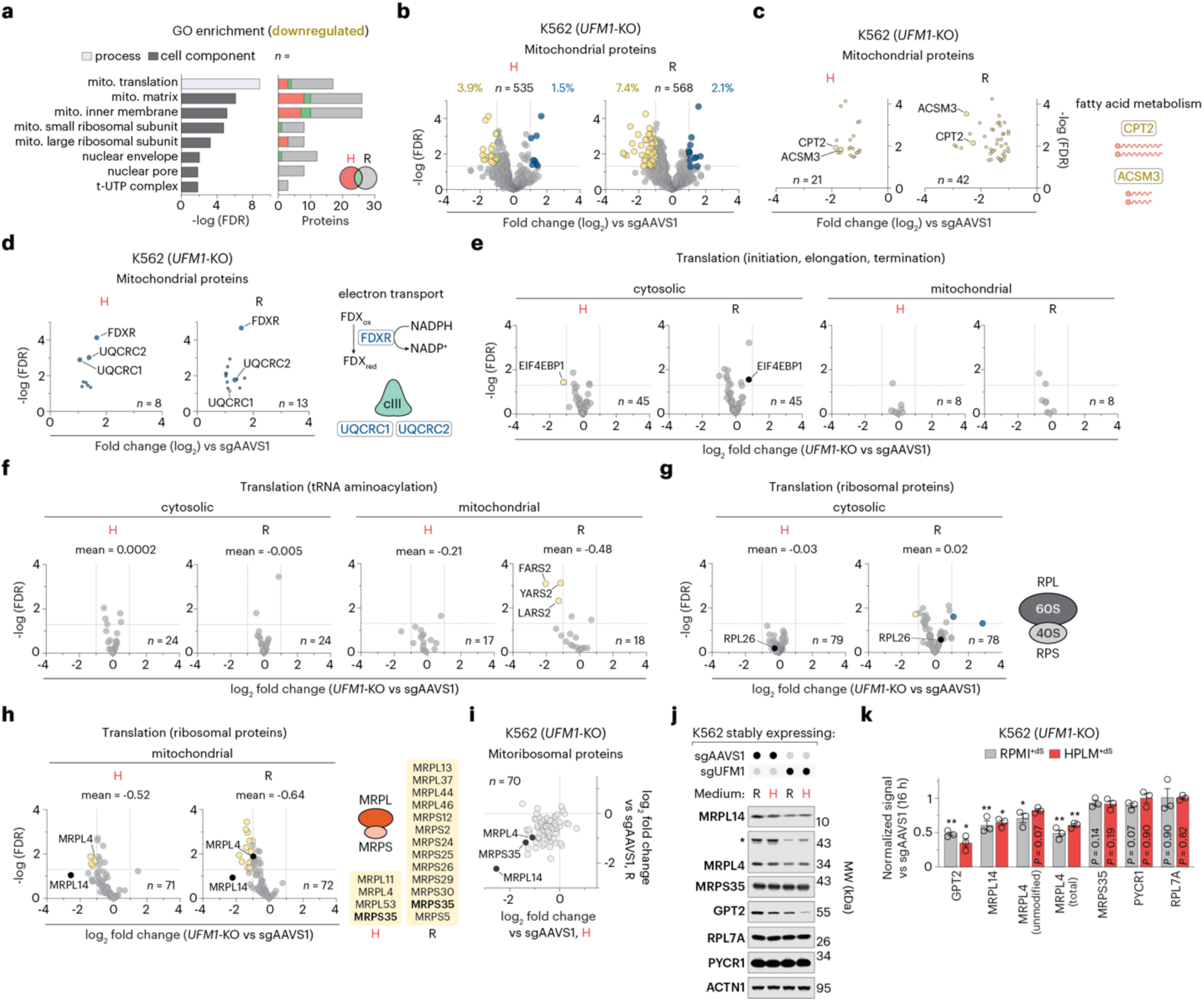
*UFM1* deletion alters mitochondrial proteostasis. (a) Enriched Gene Ontology (GO) biological processes and cellular components identified among the 309 total downregulated protein hits in *UFM1*-knockout versus control cells (left). Categories were identified using DAVID functional annotation analysis^86,87^. Number of proteins per condition in each category (right). (b-h) Relative protein levels in *UFM1*-knockout versus control cells (*n* = 6 biologically independent samples). In F-H, mean values represent the average log2 fold change across the displayed proteins. (i) Comparison of relative mitoribosomal protein levels for *UFM1*-knockout versus control cells in HPLM^+dS^ and RPMI^+dS^ (*n* = 6 biologically independent samples). (j) Immunoblots for expression of MRPL14, MRPL4, MRPS35, GPT2, RPL7A, and PYCR1 in *UFM1*-knockout and control cells. ACTN1 served as the loading control. The asterisk denotes a band corresponding to a putative monoubiquitinated MRPL4 species (∼10 kDa shift). (k) Relative protein levels quantified from immunoblot signals in *UFM1*-knockout versus control cells (mean ± s.e.m., *n* = 3 biologically independent samples). Two-tailed Welch’s *t*-test. \*\**P* < 0.01, \**P* < 0.05. For each replicate, raw signals were normalized to ACTN1 and then scaled to the maximum normalized value within that replicate. MRPL4 (unmodified) refers to the ∼35 kDa band. MRPL4 (total) represents the sum of the ∼35 kDa and ∼45 kDa signals. In b, percentages denote differentially abundant proteins among the quantified mitochondrial proteome (as defined by MitoCarta 3.0). In b-i, log2 fold change and FDR values were determined using FragPipe-Analyst^85^. In b and e-h, dotted lines denote a fold change of ± 2 (x-axis) and an FDR of 0.05 (y-axis).

Guided by these results, we examined how loss of *UFM1* affected protein abundance across the subset of over 500 mitochondrial proteins quantified, as defined by the MitoCarta 3.0 inventory^65^. We identified dozens of differentially regulated proteins within this collection and, consistent with our GO analysis, found that downregulated hits outnumbered upregulated hits by 3-fold in each condition (Fig. 7b). Among the proteins most depleted in both conditions were carnitine palmitoyltransferase 2 (CPT2) and acyl-CoA synthetase medium-chain family member 3 (ACSM3) – enzymes that facilitate fatty acid utilization (Fig. 7c). While relatively fewer hits displayed increased abundance, we found that ferredoxin reductase (FDXR) and the core Complex III subunits UQCRC1 and UQCRC2 were each upregulated regardless of culture in HPLM^+dS^ or RPMI^+dS^ (Fig. 7d). FDXR plays a critical role in electron transfer and iron-sulfur cluster biogenesis, while the two UQCRC isoforms are essential for the assembly of Complex III within the electron transport chain.

Given that mitochondrial translation was the most highly enriched category from our composite GO analysis, we systematically examined proteins involved in cytosolic and mitochondrial translation. We first observed that loss of *UFM1* had a negligible impact on initiation, elongation, and termination factors localized to either the cytosol (45 total) or mitochondria (8 total) (Fig. 7e). Similarly, *UFM1* deletion did not alter cytosolic aminoacyl-tRNA synthetase levels, whereas mitochondrial counterparts showed a modest global reduction that was exacerbated in RPMI^+dS^ (39% mean decrease; three hits) versus HPLM^+dS^ (16%; zero hits) (Fig. 7f). Next, we examined ribosomal proteins. Across nearly 80 components spanning the cytosolic large (RPL) and small (RPS) subunits, including RPL26, protein levels were virtually unaffected (Fig. 7g). By contrast, loss of *UFM1* reduced the mean abundance across roughly 70 proteins comprising the mitochondrial large (MRPL) and small (MRPS) subunits by 40% in HPLM^+dS^ (four hits) and nearly 60% (thirteen hits) in RPMI^+dS^ (Fig 7h).

To validate these results, we selected three mitoribosomal proteins for follow-up evaluation: the lone overlapping hit (MRPS35); a second downregulated in both conditions but that fell just below the hit threshold in RPMI^+dS^ (MRPL4); and a third that was most robustly depleted in each case despite not meeting the significance cutoff (MRPL14) (Fig. 7i). As anticipated, immunoblot analysis showed that *UFM1* knockout reduced MRPL14 and MRPL4 levels by 35-50% in both conditions, with GPT2 serving as a positive control for relative depletion, RPL7A as an unrelated cytosolic ribosomal control, and pyrroline-5-carboxylate reductase 1 (PYCR1) as an amino acid metabolism control (Fig. 7j, k). While immunoblotting instead indicated only a slight reduction in MRPS35 levels, this discrepancy likely stems from the inherent limitations of antibody-specific immunodetection, illustrating a sensitivity gap often encountered when validating quantitative proteomics data using this approach^66–68^. Indeed, MRPS35 exhibited a relatively low baseline abundance in our proteomics data, with intensities 60-80% lower than those of MRPL4 and MRPL14 (Extended Data Fig. 6c). Of note, we detected two distinct bands for MRPL4: one at the predicted molecular weight and a second species roughly 10 kDa heavier, consistent with the addition of a monoubiquitin (Ub) or UBL modification. Since Ub sites in MRPL4 have been previously reported^69,70^, we assessed MRPL4 levels by quantifying both the unmodified and total protein signal.

Collectively, these results demonstrate that while (de)UFMylation primarily functions at ER-localized ribosomes, this pathway exerts a broader impact on global proteostasis that extends to other subcellular compartments. Such regulation encompasses a diverse range of biological processes – including mitochondrial translation, protein transport, cellular architecture, and several branches of metabolism – and can ultimately dictate cell fitness by modulating the abundance of specific client proteins like GPT2.

### Conditional dependence on the UFMylation system varies with cell-intrinsic factors

Since *UFM1* is defined as an essential gene in roughly half of the cancer cell lines in DepMap (excluding K562)^7^, we sought to determine whether the gene-nutrient interaction between *UFM1* and alanine was conserved. Using *UFM1*-targeting and *AAVS1*-targeting sgRNAs, we transduced a panel of six blood cancer lines spanning three lines each of myeloid and lymphoid origin: K562 (CML), two acute myeloid leukemia (AML) (NOMO1 and MOLM13), one T-cell acute lymphoblastic leukemia (T-ALL) (Jurkat), a diffuse large B-cell lymphoma (DLBCL) (SUDHL4), and one anaplastic large cell lymphoma (ALCL) (Karpas299) (Fig. 8a). We then evaluated the relative growth of these *UFM1*-depleted lines in RPMI^+dS^ and either alanine-free or complete HPLM^+dS^ (Fig. 8b). Despite achieving near-complete loss of UFM1 based on immunoblot analysis, the *UFM1*-depleted K562 cell population exhibited only minor growth defects across both alanine-free conditions relative to HPLM^+dS^. Similarly, nutrient conditions had minimal impact on the relative growth of *UFM1-*depleted Jurkat, SUDHL4, and Karpas299 cells. By contrast, *UFM1* depletion in NOMO1 and MOLM13 cells led to a 25-30% greater growth impairment in RPMI^+dS^ versus HPLM^+dS^, an effect independent of alanine removal from HPLM. These results establish that *UFM1* dependence is shaped by an interplay between cell-intrinsic and extrinsic factors that extend beyond alanine availability.

**Fig. 8.**
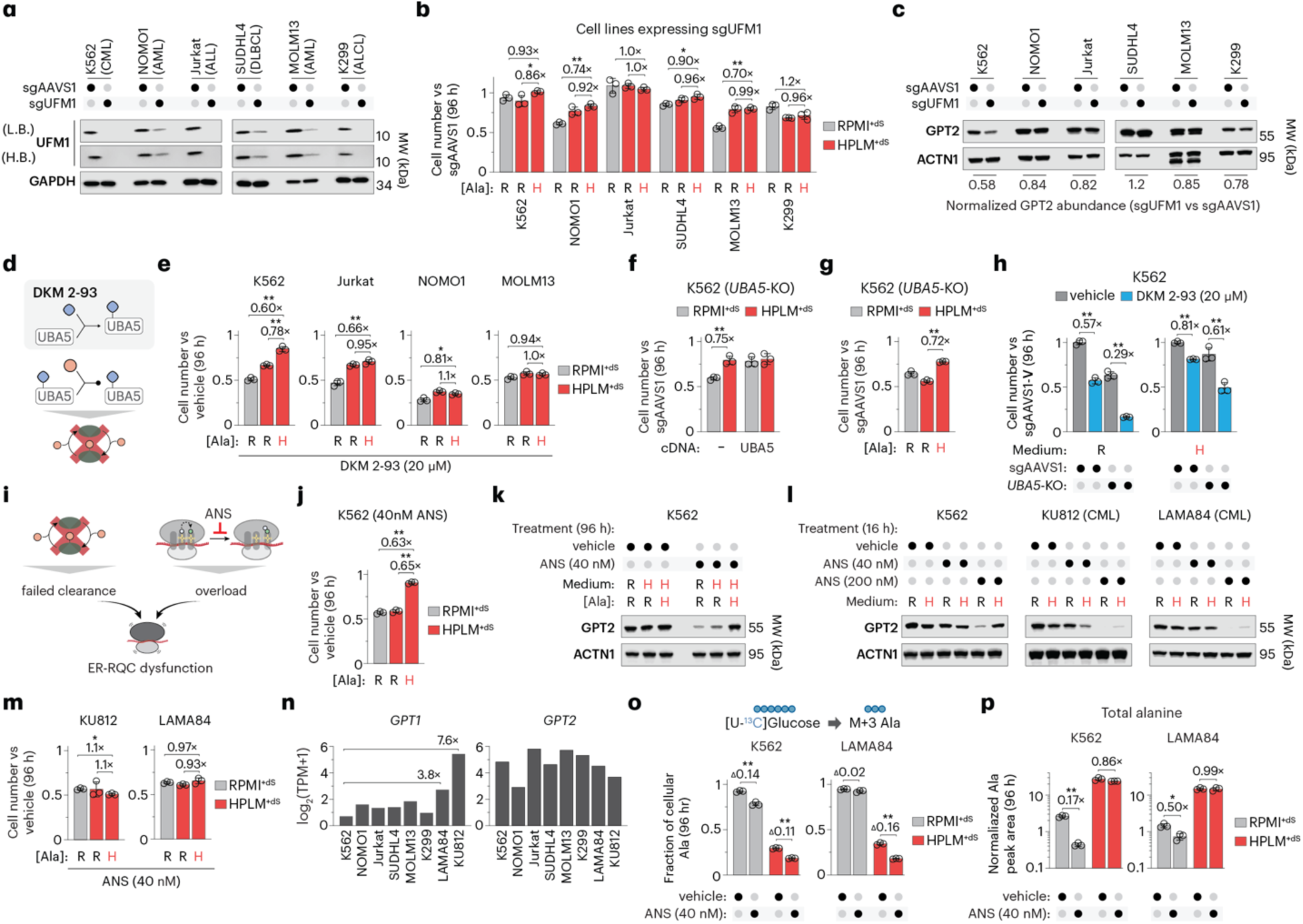
Conditional dependence on the UFMylation system varies with cell-intrinsic factors. (a, c) Immunoblots for expression of UFM1 (a) and GPT2 (c) in cell lines transduced with either sgAAVS1 or sgUFM1. GAPDH or ACTN1 served as the loading control. Bottom row (c), GPT2 signal normalized to ACTN1 in the respective cell lines expressing sgUFM1 versus sgAAVS1. L.B., low brightness. H.B., high brightness. ALCL, anaplastic large cell lymphoma; ALL, acute lymphoblastic leukemia; AML, acute myeloid leukemia; CML, chronic myeloid leukemia; DLBCL, diffuse large B cell lymphoma. (b) Relative growth of indicated cell lines transduced with sgUFM1 versus sgAAVS1 (mean ± s.d., *n* = 3 biologically independent samples). (d) Schematic of DKM 2-93, identified as a covalent ligand of UBA5 that inhibits UFM1 activation^71^. (e) Relative growth of indicated cell lines treated with DKM 2-93 versus vehicle (mean ± s.d., *n* = 3 biologically independent samples). (f, g) Relative growth of *UBA5*-knockout versus control cells (mean ± s.d., *n* = 3 biologically independent samples). (h) Relative growth of *UBA5*-knockout and control cells treated with DKM 2-93 (mean ± s.d., *n* = 3 biologically independent samples). (i) Schematic of distinct mechanisms that impair ER-RQC function. (j, m) Relative growth of indicated cell lines treated with anisomycin (ANS) versus vehicle (mean ± s.d., *n* = 3 biologically independent samples). (k, l) Immunoblots for expression of GPT2 in cell lines treated with vehicle or ANS. ACTN1 served as a loading control in both cases. (n) mRNA transcript levels for *GPT1* (left) and *GPT2* (right) from curated RNA-Seq data^26^. (o) Fractional labeling of alanine for cell lines treated with ANS versus vehicle (mean ± s.d., *n* = 3 biologically independent samples). Two-tailed Welch’s *t*-test. Values above brackets indicate differences in fractional labeling between bars. \*\**P* < 0.005. (p) Alanine levels in ANS- and vehicle-treated cells (mean ± s.d., *n* = 3 biologically independent samples). Two-tailed Welch’s *t*-test. \*\**P* < 0.005, \**P* < 0.01. In b, e-h, j, and m, Two-tailed Welch’s *t*-test comparing the respective mean ± s.d. (bar) versus mean ± s.d. (control cells) between bars. \*\**P* < 0.01, \**P* < 0.05. In b, e-h, j, m, and p, values above brackets indicate fold change between bars.

To investigate why our *UFM1*-depleted K562 population did not recapitulate the expected conditional phenotype, we assessed GPT2 levels and fractional M+3-alanine labeling from [U-^13^C]-glucose across our panel. *UFM1*-depleted K562 cells displayed a distinctly stronger reduction in GPT2 compared with the other lines, though this effect was substantially weaker than that seen in our clonal *UFM1*-knockout cells (Fig. 2g and 8c). Moreover, while *UFM1* depletion modestly decreased fractional ^13^C labeling of alanine (5-9%) across four of the six lines in HPLM^+dS^, only K562 cells showed a substantial reduction (11% versus :: 1% in the other lines) in M+3-alanine labeling during growth in RPMI^+dS^ (Extended Data Fig. 7a). Nonetheless, the magnitude of this latter defect was nearly 5-fold lower than in our clonal knockout cells. *UFM1* depletion also minimally affected protein abundance across the three mitoribosomal proteins described above, consistent with the modest impact on GPT2 in these shorter-term assays (Extended Data Fig. 7b). These results suggest that the short-term dynamics of bulk editing may mask or delay the consequences of chronic *UFM1* deficiency in some cases, potentially due to residual UFM1 or the slow turnover kinetics of specific (de)UFMylation clients. Of note, the post-selection window in our prior CRISPR screens was more than twice as long relative to the cumulative endpoint of these assays, likely providing a sufficient timeframe for downstream proteostatic effects in K562 cells to fully manifest. This shorter assay period has otherwise proven sufficient to recapitulate conditional CRISPR phenotypes for other hits from these screens^15,16^.

Given the apparent limitations of bulk *UFM1* editing in some cases, we explored chemical inhibition to acutely disrupt UFMylation with increased throughput. DKM 2-93 was previously identified as a covalent ligand of UBA5 that targets the catalytic Cys250, irreversibly blocking UFM1 activation (Fig. 8d)^71^. Treating K562 cells with this compound elicited a 40% stronger growth defect in RPMI^+dS^ versus HPLM^+dS^, though this difference was twice as large as the impairment observed in alanine-free HPLM^+dS^, suggesting these conditional responses could only be partially attributed to relative alanine availability (Fig. 8e). In addition, DKM 2-93 induced alanine-independent growth defects in NOMO1 cells, mirroring the *UFM1* depletion phenotypes noted above. By contrast, while bulk *UFM1* deficiency had little impact on Jurkat cells, DKM 2-93 elicited 30-35% stronger responses in RPMI^+dS^ regardless of alanine removal from HPLM. Moreover, while MOLM13 cells displayed a 30% greater growth defect in RPMI^+dS^ versus HPLM^+dS^ upon *UFM1* depletion, they showed no such conditional response to DKM 2-93. These results suggest that the fitness effects of DKM 2-93 likely extend beyond UBA5 inhibition.

To directly assess the specificity of DKM 2-93, we next generated clonal *UBA5*-knockout K562 cells, which displayed a 25% stronger growth defect in RPMI^+dS^ relative to HPLM^+dS^ (Fig. 8f, g and Extended Data Fig. 7c). This defect was rescued by expressing an sgRNA-resistant *UBA5* cDNA and phenocopied by removing alanine from HPLM. Consistent with additional off-target activity, while DKM 2-93 (20 µM) impaired the growth of K562 control cells to a similar extent as *UBA5* deletion, *UBA5*-knockout cells displayed further defects in HPLM^+dS^ (40%) and RPMI^+dS^ (70%) (Fig. 8h). In addition, we extended our analysis of DKM 2-93 to include two B-ALL lines (SEM and NALM6) and a non-cancer adherent line (HEK293T). These data revealed that the conditional sensitivity in SEM and HEK293T resembled that of Jurkat, whereas NALM6, similar to MOLM13, lacked a differential growth response in RPMI^+dS^ versus HPLM^+dS^ (Extended Data Fig. 7d).

Impaired (de)UFMylation prevents the clearance of ER-stalled ribosomes, while anisomycin induces ribosome collisions that saturate the ER-RQC pathway. Since both perturbations can lead to the accumulation of stalled ribosomes and ER-RQC dysfunction, we asked whether anisomycin could phenocopy conditional *UFM1* dependence in K562 cells (Fig. 8i). Notably, low-dose anisomycin (40 nM) induced a 40% greater growth defect in both RPMI^+dS^ and alanine-free HPLM^+dS^ versus complete HPLM^+dS^ (Fig. 8j). Furthermore, GPT2 levels were markedly reduced in anisomycin-treated cells across both alanine-restricted conditions, suggesting a link between GPT2 abundance and the ER-RQC response (Fig. 8k). Consistent with the translational defects observed upon loss of *UFM1*, we found that 40 nM anisomycin reduced nascent protein synthesis by 20-30% (TAMRA signal normalized to total protein) in RPMI^+dS^ and alanine-free HPLM^+dS^ but did not affect AHA incorporation in complete HPLM^+dS^ (Extended Data Fig. 7e-g). By contrast, high-dose anisomycin (200 nM) reduced TAMRA signal by over 60% in complete HPLM^+dS^, confirming that the low-dose response was exacerbated by alanine restriction. Additionally, low-dose cycloheximide (100 nM), a translation inhibitor that also binds the 60S subunit but acts through a distinct elongation-blocking mechanism, replicated the conditional sensitivity observed with anisomycin, inducing a 30% greater growth defect in both alanine-restricted conditions (Extended Data Fig. 7h, i). Together, these data support a model in which persistent ribosome stalling, regardless of the specific trigger, overwhelms the ER-RQC pathway and reduces GPT2 levels in K562 cells.

Although GPT2 abundance and M+3-alanine labeling phenotypes in our bulk-edited K562 cells were more attenuated than in our clonal knockout, these effects remained unique to K562 cells among the six lines tested. Therefore, we considered whether GPT2 might be a subtype-specific client of the ER-RQC pathway. To examine this, we evaluated GPT2 abundance following acute treatment (16 h) with low-dose (40 nM) or high-dose (200 nM) anisomycin in K562 cells and two additional CML lines with erythroid-like features: KU812 and LAMA84. In each case, GPT2 levels were modestly reduced by the low-dose treatment and severely depleted at the high dose (Fig. 8l). Surprisingly, however, KU812 and LAMA84 cells did not exhibit the same conditional growth response observed in anisomycin-treated K562 cells (Fig. 8m). Despite the near-selective expression of *GPT2* in K562 and most human cancer lines, we reasoned that GPT1 activity might compensate for GPT2 depletion in KU812 and LAMA84 cells – a rationale supported by our findings using MTS-deficient GPT2, which established that the alanine limitation is not compartmentalized. Indeed, RNA-Seq data indicate that *GPT2* is robustly expressed in all three CML lines (and in the other lines from our initial panel), whereas *GPT1* levels are 9-fold and 70-fold higher in LAMA84 and KU812, respectively, compared with K562 (Fig. 8n)^26^. To determine whether even the more modest elevation of *GPT1* in LAMA84 relative to K562 could effectively preserve de novo alanine synthesis, we assessed M+3-alanine labeling from [U-^13^C]-glucose in each of these two lines following anisomycin treatment. As expected, alanine pools from vehicle-treated cells in RPMI^+dS^ were almost completely M+3-labeled (Fig. 8o). Anisomycin reduced this fractional labeling by 14% in K562 but had a negligible impact in LAMA84, suggesting that basal GPT1 levels in the latter line are sufficient to compensate for GPT2 depletion under these conditions. By contrast, the same treatment in HPLM^+dS^ reduced fractional alanine labeling by a similar extent (11-16%) across both lines. Taken with our bulk *UFM1*-editing data, these results suggest that impaired de novo alanine synthesis is partially uncoupled from total GPT1/2 abundance in anisomycin-treated or UFMylation-deficient cells in HPLM^+dS^. Notably, anisomycin also induced a more severe depletion of the total alanine pool of K562 (80%) compared with LAMA84 (50%) in RPMI^+dS^, while having little effect on this pool for either line in HPLM^+dS^ (Fig. 8p).

Collectively, these results suggest that while the relative sensitivity of specific clients like GPT2 to ER-RQC dysfunction may partially converge by lineage, the ensuing impact on cell fitness is shaped by an interplay between extrinsic nutrient conditions and intrinsic factors, such as alanine availability and basal *GPT1* expression.

## DISCUSSION

We previously identified core components of the UFMylation system as conditionally essential for K562 leukemia cells grown in RPMI versus HPLM^12^. Here, we trace this conditional dependence to the differential availability of alanine (the second most abundant amino acid in human blood), which is provided at physiologic levels in HPLM but is absent from RPMI and other conventional synthetic media such as MEM and DMEM. We find that the UFMylation pathway maintains cellular levels of GPT2, a mitochondrial enzyme that mediates de novo alanine synthesis, although GPT2 itself is not a direct UFM1 substrate. Most human cancer lines, including K562, display near-selective expression of *GPT2* relative to *GPT1*^26^. Our study reveals that the UFMylation system serves as an ER-localized “ribosome collision counter.” By facilitating the ER-RQC-mediated clearance of stalled ribosomes, dynamic UFMylation buffers GPT2 abundance, thereby preserving cellular alanine pools required to sustain protein synthesis under alanine-restricted conditions.

Although alanine is not a defined component of RPMI, in vitro studies often rely on synthetic media supplemented with 5-20% unmodified, rather than dialyzed, serum. We previously showed that 10% FBS can provide a substantial, albeit sub-physiologic, concentration of alanine (∼100 µM) prior to dialysis^12^. Consequently, gene-nutrient interactions between the UFMylation machinery and alanine are effectively masked under typical cell culture conditions. Whether such interactions are relevant in vivo remains an open question, as systemic alanine restriction is likely unattainable due to robust de novo synthesis via the glucose-alanine (Cahill) cycle and the inter-organ buffering of plasma alanine levels^72–74^. However, localized alanine availability may be limited within poorly perfused or nutrient-depleted tumors.

Regardless of a specific in vivo context, given that translational regulation of GPT2 underlies the exogenous alanine requirement in UFMylation-deficient cells, we leveraged this finding to explore whether the system more broadly contributes to global proteostasis. Notably, *UFM1* deletion induces widespread proteomic remodeling, with differentially abundant proteins showing only modest overlap between cells cultured in HPLM and RPMI. While these client proteins span diverse biological processes, the extent of remodeling was 2-fold more extensive in RPMI, likely reflecting increased translational stress driven by relative alanine limitation. Among the most striking effects shared across both conditions was a selective depletion of mitoribosomal proteins, in contrast to cytosolic ribosome components, which were largely unaffected. Together, our work demonstrates that despite targeting a primary substrate (RPL26) on ER-localized ribosomes, the UFMylation system coordinates a multi-organelle proteostasis network downstream of ER-RQC and potentially via ER-phagy. Future studies could elucidate the specific mechanisms by which (de)UFMylation modulates the synthesis or stability of distinct clients, whether ER-resident or localized to distal compartments. Moreover, while we identify impaired translational output as the basis for conditional *UFM1* dependence, integrating ribosome profiling (Ribo-seq) with transcriptomics will be required to precisely define ribosomal occupancy and translational efficiency upon disruption of the UFMylation pathway.

Our study also demonstrates that conditional *UFM1* dependence varies across cell lines and may be dictated by differences in media composition beyond alanine. Indeed, while masking effects associated with bulk editing attenuated the metabolic consequences of *UFM1* deficiency in K562 cells, we identified two similarly edited AML cell lines that displayed increased *UFM1* dependence in RPMI versus HPLM independent of alanine availability and GPT2 abundance. Future efforts to delineate the specific gene-nutrient interactions underlying these phenotypes will be critical to further understanding *UFM1* essentiality. Nonetheless, our findings broadly suggest that the UFMylation-regulated proteome is highly context-dependent, likely varying across cancer subtypes, cell lineages, and tissues.

Given the strong co-dependency profiles between *ODR4* and core UFMylation components in DepMap, paralleling their shared identification as RPMI-essential hits in our screens, we propose that drivers of (de)UFMylation essentiality independent of the alanine-GPT2 axis are also linked to the maintenance of ER-ribosome integrity and, in turn, global proteostasis and translational output. While essentiality profiles for these components vary widely across hundreds of CRISPR screens performed in conventional media, we further propose that dependence on this system is governed by an interplay of genetic and environmental factors. Together, these determinants shape both the translational stress burden and the nature of proteomic remodeling upon impaired UFMylation, potentially giving rise to distinct, context-dependent vulnerabilities.

RNA-Seq data indicate near-universal expression of the UFMylation machinery across almost 1,700 human cancer cell lines^26^. However, since these components are also expressed in most normal tissues and are vital for organismal physiology^19,75,76^, the therapeutic window for targeting this system based solely on relative protein abundance is likely narrow. Our work suggests that identifying context-dependent proteostatic clients, or possibly additional substrates beyond RPL26, may instead reveal strategies to exploit synthetic and/or conditional lethality, thereby expanding the potential window for UFMylation-targeting therapies. The development of selective UBA5 inhibitors^77,78^ and compounds that target either the UFL1-DDRGK1-CDK5RAP3 E3 ligase complex or UFSP2^31,79,80^ will facilitate the systematic mapping of these client landscapes with greater throughput and across diverse biological models. These tools will also enable precise dissection of acute versus chronic UFMylation-dependent proteome remodeling. Indeed, while anisomycin similarly induced marked GPT2 depletion in multiple erythroid-like lines via stochastic ribosome stalling, this drug binds 60S subunits regardless of ER localization. Consequently, unlike a specific (de)UFMylation inhibitor, anisomycin not only overwhelms ER-RQC capacity but also likely elicits pleiotropic effects that extend beyond phenocopying targeted disruption of the UFMylation system.

Finally, our study suggests that the UFMylation-GPT2 axis may have broader implications for developmental biology and cell state transitions. Despite negligible overlap with our global proteomics data, Gpt2 was previously identified among a small set of proteins (< 20) depleted during the erythroid differentiation of murine precursor cells^81^. Given evidence that impaired UFMylation induces profound defects in murine erythroid development^82–84^, these observations – taken together with our findings – suggest that this system helps coordinate proteomic adaptations that drive or accompany lineage-specific transitions. Future studies will be essential to define putative links between (de)UFMylation, proteostatic buffering, and cell state across diverse biological contexts.

## Materials

| REAGENT or RESOURCE | SOURCE | IDENTIFIER |
| --- | --- | --- |
| Antibodies |  |  |
| Rabbit monoclonal anti-ACTN1 (D6F6) | Cell Signaling Technology | Cat #: 6487<br>RRID: AB_11179206 |
| Mouse monoclonal anti-Flag (M2) | Sigma-Aldrich | Cat #: F1804<br>RRID: AB_262044 |
| Mouse monoclonal anti-GAPDH (0411) | Santa Cruz Biotechnology | Cat #: sc-47724<br>RRID: AB_627678 |
| Mouse monoclonal anti-GPT2 (G-7) | Santa Cruz Biotechnology | Cat #: sc-398383<br>RRID: AB_2927429 |
| Rabbit polyclonal anti-MRPL14 | Proteintech | Cat #: 15040-1-AP<br>RRID: AB_2878102 |
| Rabbit polyclonal anti-MRPL4 | Proteintech | Cat #: 27484-1-AP<br>RRID: AB_2810294 |
| Rabbit polyclonal anti-MRPS35 | Proteintech | Cat #: 16457-1-AP<br>RRID: AB_2146521 |
| Rabbit monoclonal anti-PYCR1 (E7P3I) | Cell Signaling Technology | Cat #: 37635<br>RRID: AB_2904491 |
| Rabbit polyclonal anti-RAPTOR | EMD Millipore | Cat #: 09-217<br>RRID: AB_612103 |
| Rabbit polyclonal anti-RPL26 | Abcam | Cat #: ab59567<br>RRID: AB_945306 |
| Rabbit polyclonal anti-RPL7A (E109) | Cell Signaling Technology | Cat #: 2415<br>RRID: AB_2182059 |
| Rabbit monoclonal anti-SEC61B (D5Q1W) | Cell Signaling Technology | Cat #: 14648<br>RRID: AB_2798555 |
| Mouse monoclonal anti-TAMRA (5G5) | Thermo Fisher | Cat #: MA1-041<br>RRID: AB_2536728 |
| Rabbit polyclonal anti-UBA5 | Proteintech | Cat #: 12093-1-AP<br>RRID: AB_2211743 |
| Rabbit polyclonal anti-UFC1 | Proteintech | Cat #: 15783-1-AP<br>RRID: AB_2213938 |
| Rabbit monoclonal anti-UFL1 (F5P6L) | Cell Signaling Technology | Cat #: 96396<br>RRID: Not found |
| Rabbit polyclonal anti-UFM1 | Thermo Fisher | Cat #: PA5-58916<br>RRID: AB_2649238 |
| Mouse monoclonal anti-UFSP1 (H-3) | Santa Cruz Biotechnology | Cat #: sc-398577<br>RRID: Not found |
| Mouse monoclonal anti-UFSP2 (G-11) | Santa Cruz Biotechnology | Cat #: sc-376084<br>RRID: AB_10989729 |
| Rabbit monoclonal anti-Mouse IgG (Light Chain Specific) HRP (D3V2A) | Cell Signaling Technology | Cat #: 58802<br>RRID: AB_2799549 |
| Mouse monoclonal anti-Rabbit IgG (Conformation specific) (L27A9) | Cell Signaling Technology | Cat #: 3678<br>RRID: AB_1549606 |
| Horse anti-Mouse IgG HRP | Cell Signaling Technology | Cat #: 7076<br>RRID: AB_330924 |
| Goat anti-Rabbit IgG HRP | Cell Signaling Technology | Cat #: 7074<br>RRID: AB_2099233 |
| Mouse monoclonal IgG2a isotype control | Cell Signaling Technology | Cat #: 61656<br>RRID: AB_2799613 |
| Chemicals and commercial kits |  |  |
| 3X FLAG Peptide | Sigma-Aldrich | Cat #: F4799 |
| Anisomycin | Selleck | Cat #: S7409 |
| Anti-FLAG M2 Affinity Gel | Sigma-Aldrich | Cat #: A2220 |
| Blasticidin (solution) | Invivogen | Cat #: ant-bl-1 |
| Chloroform | Thermo Fisher | Cat #: 032614.K2 |
| Click-IT AHA (L-Azidohomoalanine) | Thermo Fisher | Cat #: C10102 |
| Cycloheximide | Selleck | Cat #: S7418 |
| D-Glucose (U-<sup>13</sup>C<sub>6</sub>, 99%) | Cambridge Isotope Labs | Cat #: CLM-1396 |
| Digitonin | Sigma-Aldrich | Cat #: D141 |
| DMEM, high glucose, GlutaMAX | Thermo Fisher | Cat #: 10566024 |
| Fetal Bovine Serum (FBS), Heat Inactivated | Thermo Fisher | Cat #: 16140071 |
| Glucose | Thermo Fisher | Cat #: 15023021 |
| L-Alanine | Sigma-Aldrich | Cat #: A7627 |
| Mixture of amino acid standards for metabolomics | Cambridge Isotope Labs | Cat #: MSK-A2-1.2 |
| Penicillin-Streptomycin | Thermo Fisher | Cat #: 15140122 |
| PowerUp SYBR Green master mix | Thermo Fisher | Cat #: A25742 |
| Puromycin dihydrochloride | Sigma-Aldrich | Cat #: P7255 |
| QIAzol lysis reagent | QIAGEN | Cat #: 79306 |
| RPMI 1640 | Thermo Fisher | Cat #: 11875093 |
| RPMI 1640, no glucose | Thermo Fisher | Cat #: 11879020 |
| SimplyBlue SafeStain | Thermo Fisher | Cat #: LC6060 |
| Sucrose | Sigma-Aldrich | Cat #: 84097 |
| SUPERase-In RNase Inhibitor | Thermo Fisher | Cat #: AM2694 |
| SuperScript IV First-Strand Synthesis System | Thermo Fisher | Cat #: 18091050 |
| Synthetic medium components | Multiple | See Table S1 |
| TAMRA Alkyne | Thermo Fisher | Cat #: T10183 |
| Trichloroacetic acid | Sigma-Aldrich | Cat #: T6399 |
| X-tremeGENE 9 DNA Transfection Reagent | Sigma-Aldrich | Cat #: 6365779001 |
| Click-IT Protein reaction buffer kit | Thermo Fisher | Cat #: C10276 |
| miRNeasy mini kit | QIAGEN | Cat #: 217004 |
| Deposited Data |  |  |
| Differential dependency data, genome-wide CRISPR screens in K562 cells | Rossiter et al., 2021 | Supp. Table 1 |
| DepMap gene dependency data, core and accessory UFMylation components | DepMap 24Q4 Public | https://depmap.org/portal/data_page/?tab=allData |
| Transcript level data, core UFMylation components, GPT1, and GPT2 in cancer lines | DepMap 24Q4 Public | https://depmap.org/portal/data_page/?tab=allData |
| Flag-based affinity purification prey proteins for wild-type UFM1, UFM1ΔC3, and RAP2A | This paper | Supp. Table 3 and TBD |
| Label-free quantitative proteomics | This paper | Supp. Table 4 and TBD |
| Experimental Models: Cell Lines |  |  |
| K562 | J.D. Griffin | RRID_CVCL_0004 |
| NOMO1 | J.D. Griffin | RRID_CVCL_1609 |
| MOLM-13 | CCLE | RRID_CVCL_2119 |
| SUDHL4 | M.A. Shipp | RRID_CVCL_0539 |
| Jurkat | ATCC | RRID_CVCL_0065 |
| Karpas299 | D.M. Sabatini (lab cell bank) | RRID_CVCL_1324 |
| SEM | CCLE | RRID_CVCL_0095 |
| LAMA84 | CCLE | RRID_CVCL_0388 |
| NALM6 | D.M. Weinstock | RRID_CVCL_0092 |
| KU812 | ATCC | RRID_CVCL_0379 |
| Oligonucleotides |  |  |
| Primers for construction of expression plasmids | This paper | Supp. Table 5 |
| Primers for qPCR (GPT2, ACTB) | This paper | Supp. Table 5 |
| Individual sgRNA target sequences | This paper | Supp. Table 5 |
| gBlock sequences (UBA5, UFM1, UFSP1, UFSP2) | This paper | Supp. Table 5 |
| Recombinant DNA |  |  |
| pLJC6-Rap2A-3xFLAG | Rossiter et al.,2021 | Addgene 163445 |
| pLJC6-EV | Rossiter et al.,2021 | Addgene 136454 |
| pLJC6-GPT2-3xFLAG | Rossiter et al.,2021 | Addgene 163449 |
| pLJC6-GPT2 (ΔMTS)-3xFLAG | This paper | Addgene TBD |
| pLJC6-3xFLAG-UFM1 | This paper | Addgene TBD |
| pLJC6-3xFLAG-UFM1(ΔC2) | This paper | Addgene TBD |
| pLJC6-3xFLAG-UFM1(ΔC3) | This paper | Addgene TBD |
| pLJC6-UBA5-3xFLAG | This paper | Addgene TBD |
| pLJC6-mCherry-eGFP-RAMP4 | This paper | Addgene TBD |
| pLJC2-Blast-Rap2A-3xFLAG | Flickinger et al., 2025 | Addgene 233890 |
| pLJC2-Blast-UFSP1 | This paper | Addgene TBD |
| pLJC2-Blast-UFSP2 | This paper | Addgene TBD |
| pLJC2-Blast-UFSP2 (C302S) | This paper | Addgene TBD |
| pLentiCRISPR-v1 | Addgene | Addgene 49535 |
| pLentiCRISPR-v1-sgAAVS1 | Wang et al., 2015 | Addgene 70661 |
| pLentiCRISPR-v1-UFM1 | This paper | Addgene TBD |
| pLentiCRISPR-v1-UFSP2 | This paper | Addgene TBD |
| pLentiCRISPR-v1-UBA5 | This paper | Addgene TBD |
| TetOn-mCherry-eGFP-RAMP4 | Liang et al., 2018 | Addgene 109014 |

## Methods

### Cell lines

The Jurkat and KU-812 cell lines were purchased from ATCC. The following cell lines were kindly provided by: K562 and NOMO1, J. Griffin (Dana Farber Cancer Institute); SUDHL4, M.A. Shipp (Dana Farber Cancer Institute); NALM6, D.M. Weinstock (Dana Farber Cancer Institute); Karpas299, D.M. Sabatini (Institute of Organic Chemistry and Biochemistry of the Czech Academy of Sciences; from lab cell bank); MOLM-13, SEM, and LAMA84, Cancer Cell Line Encyclopedia (Broad Institute). Cell lines were verified to be free of mycoplasma contamination and their identities were authenticated by short tandem repeat (STR) profiling.

### Cell culture conditions

The following culture media were used (all contained 0.5% penicillin-streptomycin):

1. RPMI^+S^: RPMI 1640, no glucose (Thermo Fisher) with 5 mM glucose and 10% FBS.
2. RPMI^+dS^: RPMI 1640, no glucose (Thermo Fisher) with 5 mM glucose and 10% dialyzed FBS.
3. RPMI11^+S^: RPMI1640 (Thermo Fisher) with 10% FBS.
4. RPMI11^+2S^: RPMI1640 (Thermo Fisher) with 20% FBS.
5. DMEM^+S^: DMEM, high glucose, GlutaMAX (Thermo Fisher) with 10% FBS.
6. DMEM^+2S^: DMEM, high glucose, GlutaMAX (Thermo Fisher) with 20% FBS.
7. HPLM^+dS^: manually prepared HPLM (Supplementary Table 1) with 10% dialyzed FBS.
8. RPMI-AHA (Supplementary Table 1), no FBS.
9. HPLM-AHA (Supplementary Table 1), no FBS.

Using SnakeSkin tubing (Thermo Fisher PI88244), FBS was dialyzed as previously reported^14^. Before use, all FBS-supplemented media were sterile-filtered using a bottle-top vacuum filter with cellulose acetate membrane, pore size 0.2 μm (Corning 430626 or Nalgene 290-4520). Cells were maintained at 37°C, atmospheric oxygen, and 5% CO2.

#### Construction of knockout plasmids

For each of the following genes, sense and antisense oligonucleotides for a single guide RNA (sgRNA) targeting the respective gene were annealed and then cloned into *BsmBI*-digested pLentiCRISPR-v1: *UFM1*, *UFSP2*, and *UBA5*.

#### Construction of expression plasmids

The following genes were amplified from codon-optimized gBlock Gene Fragments (IDT), digested with *PacI-NotI*, and then cloned into pLJC6-Rap2A-3xFLAG: (1) 3xFLAG-UFM1 using primers UFM1-F/UFM1-R; (2) UBA5-3xFLAG using primers UBA5-F/UBA5-R. UFM1-F was designed to incorporate an N-terminal 3xFLAG tag and GGS linker. UFM1-R was designed to incorporate two consecutive stop codons preceding the *NotI* site. The following cDNAs were amplified from pLJC6-3xFLAG-UFM1, digested with *PacI-NotI*, and then cloned into pLJC6-Rap2A-3xFLAG: (1) 3xFLAG-UFM1(ΔC2) using primers LJC-F/UFM1_d2-R; (2) 3xFLAG-UFM1(ΔC3) using primers LJC-F/UFM1_d3-R.

The following genes were amplified from codon-optimized gBlock Gene Fragments (IDT), digested with *PacI-NotI*, and then cloned into pLJC2-Blast-Rap2A-3xFLAG: (1) UFSP1 using primers UFSP1-F/UFSP1-R; (2) UFSP2 using primers UFSP2-F/UFSP2-R. UFSP1-R and UFSP2-R were each designed to incorporate two consecutive stop codons preceding the *NotI* site. The plasmid pLJC2-Blast-UFSP2 (C302S) was generated using a 2-step protocol based on overlap extension PCR. In the first step, two fragments were amplified from pLJC2-Blast-UFSP2 using the following primer pairs: LJC2-F/UFSP2_C302S-R and UFSP2_C302S-F/LJC-R. In the second step, the two fragments were pooled in a PCR containing primers LJC2-F/LJC-R, digested with *PacI-NotI*, and then cloned into pLJC2-Blast-RAP2A-3xFLAG.

The GPT2 (ΔMTS)-3xFLAG cDNA was amplified from pLJC6-GPT2-3xFLAG using primers GPT2_MTS-F/LJC-R, digested with *PacI-NotI*, and cloned into pLJC6-Rap2A-3xFLAG. GPT2_MTS-F was designed to introduce an initiating Met immediately upstream of Ser25, based on a predicted GPT2-MTS cleavage site at residue Gln24^88^. The mCherry-eGFP-RAMP4 cDNA was amplified from TetOn-mCherry-eGFP-RAMP4 using primers EATR-F/EATR-R, digested with *PacI-NotI*, and cloned into pLJC6-Rap2A-3xFLAG. EATR-R was designed to incorporate two consecutive stop codons preceding the *NotI* site.

#### Lentivirus production

To produce lentivirus, HEK293T cells in DMEM^+S^ were co-transfected with VSV-G envelope plasmid, Delta-VPR packaging plasmid, and a transfer plasmid (pLJC6, pLJC2-Blast, or pLentiCRISPR-v1 backbone) using X-tremeGENE 9 Transfection Reagent (Sigma-Aldrich). Medium was exchanged with fresh DMEM^+2S^ 16 h after transfection, and the virus-containing supernatant was collected at 48 h post-transfection, passed through a 0.45 μm filter to eliminate cells, and then stored at -80°C.

#### Cell line construction

##### Genetic knockout clonal cell lines

To establish knockout clonal lines (*UFM1*, *UFSP2*, and *UBA5*), K562 cells were seeded at a density of 500,000 cells/mL in a 6-well plate containing 2 mL RPMI11^+S^, 8 μg/mL polybrene, and the respective pLentiCRISPR-v1 lentivirus of interest. Spin infection was carried out by centrifugation at 1,000 *g* for 45 min at 37°C. Following 16-18 h incubation, cells were pelleted to remove virus and then re-seeded into fresh RPMI11^+S^ for 24 h. Cells were then pelleted and re-seeded into RPMI11^+S^ containing puromycin for 72 h. Following selection, the population was single-cell FACS-sorted into 96-well plates containing RPMI11^+2S^ (BD FACSMelody Cell Sorter). After 1.5-2 weeks, cell clones with the desired knockout were identified by immunoblotting. To control for infection, a population of K562 cells was similarly selected following transduction with sgAAVS1-containing lentivirus.

##### UFM1-knockout cell lines

The procedure to establish knockout cell lines for short-term growth assays and other experiments was similar to that described for the clonal cell line except that cells were not FACS-sorted following puromycin selection (5 days post infection) and other minor modifications (See **Short-term growth assays**, **Acute protein level analysis**, and **Metabolite profiling and quantification of metabolite abundance**; Cells: glucose labeling).

1. Cells were transduced in parallel with either sgUFM1- or sgAAVS1-containing lentivirus.
2. The following cell lines were seeded at a density of 500,000 cells/mL: K562, Karpas299, and MOLM-13.
3. The following cell lines were seeded at a density of 1 million cells/mL: Jurkat, NOMO1, and SUDHL4.

##### cDNA expression cell lines

To establish stable expression cell lines, *UFM1*-knockout, *UFSP2*-knockout, or *UBA5*-knockout K562 clonal cells were seeded in 6-well plates containing 2 mL of RPMI11^+S^, 8 μg/mL polybrene, and either the pLJC6 or pLJC2-Blast lentivirus of interest. Spin infection and medium exchange were performed identically as described above for the knockout cell lines. Cells were then pelleted and resuspended into fresh RPMI11^+S^ containing blasticidin for 72 h. Stable cDNA expression was confirmed by immunoblotting in all cases, except for mCherry-eGFP-RAMP4, which was verified by ratiometric flow cytometry.

#### Cell lysis for immunoblotting

Cells were centrifuged at 250 *g* for 5 min, resuspended in 1 mL ice-cold PBS, and then centrifuged again at 250 *g* for 5 min at 4°C. Cells were then immediately lysed with ice-cold lysis buffer (40 mM Tris-HCl pH 7.4, 1% Triton X-100, 100 mM NaCl, 5 mM MgCl2, and 1 tablet of EDTA-free protease inhibitor (Roche 11580800; per 25-mL buffer)) unless otherwise noted. Cell lysates were cleared by centrifugation at 21,130 *g* for 10 min at 4°C and quantified for protein concentration using an albumin standard (Thermo Fisher 23209) and Bradford reagent (Bio-Rad 5000006). Clarified lysates were normalized for protein content, denatured upon the addition of 5x SDS-PAGE sample buffer (Thermo Fisher 39000), resolved by SDS-PAGE, and then transferred to a polyvinyl difluoride membrane (Millipore IPVH07850). Membranes were blocked with 5% nonfat dry milk in Tris Buffered Saline with Tween (TBST) for 1 h at room temperature, and then incubated with primary antibodies in 5% nonfat dry milk in TBST overnight at 4°C. Primary antibodies to the following proteins were used at the indicated dilutions: ACTN1 (1:2,000), FLAG (1:750), GAPDH (1:1,000), GPT2 (1:100), MRPL14 (1:500), MRPL4 (1:10,000), MRPS35 (1:10,000), PYCR1 (1:2,000), RAPTOR (1:1,000), RPL26 (1:2,500), RPL7A (1:1,000), SEC61B (1:2,000), TAMRA (1:1,000), UBA5 (1:500), UFC1 (1:500), UFL1 (1:1,000), UFM1 (1:500), UFSP1 (1:100), and UFSP2 (1:200).

Membranes were washed with TBST three times for 5 min each and then incubated with species-specific secondary antibody in 5% nonfat dry milk for 1 h at room temperature. Secondary antibodies were used at the indicated dilutions: anti-Mouse IgG (Light Chain Specific) HRP (1:1,000) (used only for the GPT2 immunoblot in Fig. 2l), anti-Rabbit IgG (Conformation specific) (L27A9) (1:2,000) (used only for the immunoblots in Fig. 3l), anti-Mouse IgG HRP (1:3,000), and anti-Rabbit IgG HRP (1:3,000). Membranes were washed again with TBST three times for 5 min each and then visualized with chemiluminescent substrate (Thermo Fisher) on a LI-COR Odyssey FC, except for the membranes in Fig. 3l, which were instead incubated with anti-Mouse IgG HRP for 1 h at room temperature prior to analogous washes and imaging.

### Short-term growth assays

#### Engineered K562 cell lines

Following at least two passages in RPMI^+S^, cells were pelleted and resuspended to a density of 100,000 cells/mL in either HPLM^+dS^ or RPMI^+dS^. After a 72-h incubation, cells were pelleted and resuspended to 1 million cells/mL in the respective parent medium. From each resuspension, cells were seeded at a final density of 20,000 cells/mL into each of three replicate wells of a 6-well plate containing 4 mL of the appropriate medium. After 96-h incubation, cell density measurements were recorded using a Coulter Counter (Beckman Z2 or M4E) with a diameter setting of 8-30 μm.

The short-term growth assay procedure using different media derivatives was identical to that above with minor modifications:

1. Modified concentrations of amino acids or salts in HPLM were incorporated at the preceding medium-specific 72-h passage step. Removal of HPLM-specific metabolites from HPLM was similarly introduced at the 72-h passage step. For stock solutions of amino acids, salts, and HPLM-specific metabolites, see Supplementary Table 1.
2. Alanine was either removed from or serially diluted in HPLM, or added to RPMI, starting at the preceding medium-specific 72-h passage step. For the amino acid dropout assays (Fig. 5), specific amino acids – including alanine – were removed from HPLM only for the final 96-h incubation, with the preceding 72-h step carried out in HPLM^+dS^.

#### Drug treatments

The short-term growth assay procedure was identical to that above with minor modifications:

1. Following 1-h incubation of seeded 6-well plates, compounds were added at specified doses and then plates were gently shaken for 2 min before the final 96-h incubation step.
2. Stock solutions were prepared as follows: anisomycin (40 mM in DMSO), cycloheximide (40 mM in water), and DKM 2-93 (100 mM in DMSO). All wells, including the untreated controls, contained 0.25% DMSO for the anisomycin and DKM 2-93 assays.
3. The following cell lines were seeded at a density of 20,000 cells/mL for the 96-h incubation step: K562, NOMO1, MOLM13, LAMA84, and NALM6. The SEM and KU812 lines were seeded at a density of 40,000 cells/mL, and the Jurkat line at 50,000 cells/mL. For the HEK293T line, 10,000 cells per well were seeded 24 h before the addition of DKM 2-93.

#### Panel of UFM1-knockout cell lines

To generate knockout populations, cell lines were transduced in parallel with sgUFM1- or sgAAVS1-containing lentivirus, and FACS sorting was not performed following puromycin selection. Instead, the cells were pelleted and resuspended to a density of 250,000 cells/mL in 12 mL of RPMI^+S^. After 48 h incubation (7 days post infection), cells were pelleted and resuspended to a density of 300,000 cells/mL in 12 mL of either HPLM^+dS^ or RPMI^+dS^. After 48 h incubation (9 days post infection), pools of cells were pelleted and seeded at a density of either 20,000 cells/mL (K562, Karpas299, NOMO1, MOLM-13, and SUDHL4) or 40,000 cells/mL (Jurkat) in each of three replicate wells in 6-well plates containing either 4 mL of the parent medium or 4 mL of the parent medium prepared with 5 mM [U-^13^C]-glucose. HPLM-resuspended cells were used to seed the wells containing either alanine-free or complete HPLM^+dS^. RPMI-resuspended cells were further used to seed T-25 culture flasks containing 12 mL RPMI^+dS^ at these same final densities. In parallel, pools of cells were pelleted and seeded at a density of 200,000 cells/mL in T-75 flasks containing 20 mL of the parent medium.

1. For 6-well plate cultures in media with unlabeled glucose, cell density measurements were recorded after 96 h incubation per the short-term growth assay procedure described above (13 days post infection).
2. For 6-well plate cultures in media prepared with [U-^13^C]-glucose, see **Metabolite profiling and quantification of metabolite abundance;** Cells: glucose labeling).
3. For the T-25 flask cultures, whole-cell lysates were generated after 96 h incubation and used for the immunoblots in Fig. 8a, c (see **Cell lysis for immunoblotting**). The relative depletion of *GPT2* in each cell line was assessed by immunoblotting, whereby GPT2 signal was normalized by the ACTN1 signal in the respective cell lines expressing sgUFM1 versus sgAAVS1 (LI-COR Image Studio 5.2).
4. For the T-75 flask cultures, see **Acute protein level analysis.**

### Acute protein level analysis

#### Engineered K562 and wild-type cell lines

Following at least two passages in RPMI^+S^, cells were pelleted and seeded at a density of 200,000 cells/mL in either HPLM^+dS^ or RPMI^+dS^. After 48 h incubation, cells were pelleted and re-seeded to that same initial density in the respective parent medium. After 48 h incubation, cells were pelleted and resuspended to a final density of 250,000 cells/mL in T-75 culture flasks containing 20 mL of the appropriate medium. After 16-18 h incubation, whole-cell lysates were generated (see **Cell lysis for immunoblotting**).

#### Drug treatments

The acute protein level analysis procedure was identical to that above with minor modifications:

1. Cells were pelleted and resuspended to a final density of 250,000 cells/mL in T-75 culture flasks containing 20 mL of the appropriate medium with either vehicle (DMSO) or anisomycin (40 nM or 200 nM).
2. The stock solution of anisomycin was prepared at (40 mM in DMSO). All flasks, including the untreated controls, contained 0.1% DMSO.

#### Panel of UFM1-knockout cell lines

See **Short-term growth assays**. At 9 days post infection, pools of transduced cells were pelleted and seeded to a density of 200,000 cells/mL in T-75 flasks containing 20 mL of the parent medium (HPLM^+dS^ or RPMI^+dS^). After 48 h incubation, cells were pelleted and resuspended to a final density of 250,000 cells/mL in T-75 culture flasks containing 20 mL of the appropriate medium. After 16-18 h incubation, whole-cell lysates were generated and used for the immunoblots in Extended Data Fig. 7 (12 days post infection) (see **Cell lysis for immunoblotting**).

### Metabolite profiling and quantification of metabolite abundance

Liquid chromatography-mass spectrometry (LC-MS) analyses were performed on a QExactive HF benchtop orbitrap mass spectrometer equipped with an Ion Max API source and HESI II probe, coupled to a Vanquish Horizon UHPLC system (Thermo). External mass calibration was performed using positive and negative polarity standard calibration mixtures every 7 days. Acetonitrile was hypergrade for LC-MS (Millipore Sigma), and all other solvents were Optima LC-MS grade (Thermo).

#### Engineered K562 cell lines

Following at least two passages in RPMI^+S^, cells were pelleted and resuspended to a density of 250,000 cells/mL in either HPLM^+dS^ or RPMI^+dS^. After 48-hr incubation, 2-3 million cells were pelleted and resuspended to a density of 1 million cells/mL of the respective parent medium. From each resuspension, cells were seeded at a final density of 20,000 cells/mL in each of three replicate wells in 6-well plates containing 4 mL of appropriate medium prepared with 5 mM of [U-^13^C]-glucose. After 96-h incubation, a 500 μL aliquot from each well was used to measure cell number and volume via Coulter Counter (Beckman) with a diameter setting of 8-30 μm. The remaining cells were centrifuged at 250*g* for 5 min, resuspended in 1 mL of ice-cold 0.9% sterile NaCl (Growcells, MSDW1000), and centrifuged at 250 *g* for 5 min at 4°C. Metabolites were extracted in 500 μL of ice-cold 80% methanol containing 500 nM internal amino acid standards (Cambridge Isotope Laboratories). Following a 10-min vortex step and centrifugation at 21,130 *g* for 10 min at 4°C, 2.5 μL from each sample was injected onto a ZIC-pHILIC 2.1 x 150 mm analytical column equipped with a 2.1 x 20 mm guard column (both were 5-μm particle size, Millipore Sigma). Buffer A was 20 mM ammonium carbonate and 40 mM ammonium hydroxide; buffer B was acetonitrile. The chromatographic gradient was run at a flow rate of 0.15 mL/min as follows: 0-20 min: linear gradient from 80 to 20% B; 20.5-28 min: hold at 80% B. The mass spectrometer was operated in full-scan, polarity-switching mode with the spray voltage set to 3.0 kV, the heated capillary held at 275°C, and the HESI probe held at 350°C. The sheath gas flow rate was set to 40 units, the auxiliary gas flow was set to 15 units, and the sweep gas flow was set to 1 unit. The MS data acquisition in positive mode was performed in a range of 50 to 750 mass/charge ratio (m/z), with the resolution set to 120,000, the Automated Gain Control (AGC) target at 10^6^, and the maximum integration time at 20 msec. The settings in negative mode were the same except that the range was instead 70 to 1000 m/z.

For the highly targeted analysis of pyruvate isotopologues, targeted selected ion monitoring (tSIM) scans were added with the following settings: resolution set to 120,000, an AGC target of 10^5^, maximum integration time of 200 msec, and isolation window of 1.0 m/z. These scans were run in negative mode for the following the target masses: 87.0088 (unlabeled), 88.0121 (M+1-labeled pyruvate), 89.0155 (M+2-labeled pyruvate), and 90.0189 (M+3-labeled pyruvate).

#### Anisomycin-treated cells

Following at least two passages in RPMI^+S^, cells were pelleted and resuspended to a density of 250,000 cells/mL in either HPLM^+dS^ or RPMI^+dS^. After 48-hr incubation, 3 million cells were pelleted and resuspended to a density of 1 million cells/mL of the respective parent culture medium. From each resuspension, cells were seeded at a final density of 20,000 cells/mL in each of three replicate wells in 6-well plates containing 4 mL of appropriate culture medium prepared with 5 mM [U-^13^C]-glucose. Following 1-h incubation of seeded 6-well plates, anisomycin (40 nM) was added and then plates were gently shaken for 2 min. After 96-h incubation, metabolite extractions were performed identically to those described above.

#### Panel of UFM1-knockout cell lines

See **Short-term growth assays.** At 9 days post-infection, pools of transduced cells were pelleted and seeded at a final density of 20,000 cells/mL (K562, Karpas299, NOMO1, MOLM-13, and SUDHL4) or 40,000 cells/mL (Jurkat) in each of three replicate wells in 6-well plates containing 4 mL of the appropriate medium with 5 mM [U-^13^C]-glucose. After 96 h incubation, metabolite extractions were performed identically to those described above.

#### Identification and quantification

Metabolite identification and quantification were performed with XCalibur v 4.1 (Thermo) using a 10-ppm mass accuracy window and 0.5 min retention time window. To confirm metabolite identities, a manually constructed library of chemical standards was used. Standards were validated by LC-MS to confirm that they generated robust peaks at the expected m/z ratio. Since total amino acid pools were assessed from cells grown in media with 5 mM [U-^13^C]-glucose, peak areas corresponding to the unlabeled species and the appropriate ^13^C isotopologues were evaluated in each case (see Supplementary Table 2), and in turn, natural isotope abundance corrections of isotopologues were performed using the AccuCor code^89^ in R studio.

### Immunoprecipitation of endogenous GPT2

Following at least two passages in RPMI^+S^, cells were pelleted and seeded at a density of 200,000 cells/mL in either HPLM^+dS^ or RPMI^+dS^. After 48-h incubation, cells were pelleted and seeded at a final density of 150,000 cells/mL in T-75 culture flasks containing 30 mL of the appropriate medium. Following an additional 72 h-incubation, cells were pelleted, rinsed once in 1 mL ice-cold PBS, and then immediately lysed in ice-cold lysis buffer (see **Cell lysis for immunoblotting**). Cell lysates were cleared by centrifugation at 21,130 *g* for 10 min at 4°C and quantified for protein concentration using an albumin standard (Thermo Fisher 23209) and Bradford reagent (Bio-Rad 5000006).

From each clarified lysate, 800 μg protein was mixed with 2 μg of either anti-GPT2 (Santa Cruz Biotechnology sc-398383) or an IgG2a isotype control (Cell Signaling Technology 61656) in a total volume of 200 μL, then incubated with rotation overnight at 4°C. For immunoprecipitation, protein A magnetic beads (Cell Signaling Technology 73778) were washed twice in lysis buffer and then 25 μL of the washed beads were added to each sample for 1 h incubation with rotation at 4°C. Beads were magnetically isolated (Thermo Fisher, DynaMag 12320D) and washed five times with 1 mL ice-cold lysis buffer. After the final wash step, samples were denatured upon addition of 75 μL 5x SDS-PAGE sample buffer (Thermo Fisher 39000). For immunoblot analysis, 10 μg of whole-cell lysate and 10 μL of corresponding protein A eluent were resolved by SDS-PAGE.

### Sucrose gradient sedimentation for polysome profiling

#### Engineered K562 cell lines

Following at least two passages in RPMI^+S^, cells were pelleted and seeded at a density of 200,000 cells/mL in either HPLM^+dS^ or RPMI^+dS^. After 48 h incubation, cells were pelleted and re-seeded at this same density in the respective parent medium. After 48 h incubation, cells were pelleted and seeded at a density of 250,000 cells/mL in T-75 culture flasks containing 45 mL of the appropriate medium. After 16-18 h incubation, cells were washed twice with 1 mL ice-cold PBS containing 100 μg/mL cycloheximide and then immediately lysed in 2.5 mL ice-cold polysome buffer (20 mM Tris-HCl pH 7.4, 1% Triton X-100, 100 mM NaCl, 10 mM MgCl2, 1mM DTT, 1 tablet of EDTA-free protease inhibitor (per 25-mL buffer), 10U/mL SUPERase·In RNase Inhibitor, and 100 μg/mL cycloheximide). Cell lysates were cleared by centrifugation at 13,000 *g* for 10 min at 4°C, transferred to a new 1.5 mL tube, and then quantified for RNA concentration (A260) using a NanoDrop One (Thermo Fisher).

Sucrose gradient preparation and fractionation were performed as previously described^90^. In brief, lysates were layered onto a 10-50% (w/v) sucrose gradient containing 20 mM Tris-HCl pH 7.4, 100 mM NaCl, 10 mM MgCl2, 10U/mL SUPERase·In RNase Inhibitor, and 100 μg/mL cycloheximide. Samples were centrifuged at 222,227 *g* in an SW41Ti rotor at 4°C for 3 h. Gradients were fractionated using a piston fractionator (BioComp), and the A260 profile was continuously monitored with a Triax UV detector and flow cell. Collected fractions were stored at -80°C until further processing.

To quantify the abundance of ribosomal subunits (40S, 60S) and monosomes (80S), the A260 profiles were analyzed in GraphPad Prism. The area-under-the-curve (AUC) for peaks corresponding to individual subunits or monosomes was determined by baseline-subtraction and integration and these values were then normalized to the input A260 of the respective sample.

#### Anisomycin-treated cells

The polysome profiling procedure was identical to that above with minor modifications:

1. Cells were pelleted and resuspended to a final density of 250,000 cells/mL in T-75 culture flasks containing 45 mL of the appropriate medium with either vehicle (DMSO) or anisomycin (40 nM or 200 nM).
2. The stock solution of anisomycin was prepared at (40 mM in DMSO). All flasks, including the untreated controls, contained 0.1% DMSO.

For immunoblotting of gradient fractions, proteins were precipitated by adding 20% TCA (v/v) for 10 min at 4°C. Precipitates were collected by centrifugation at 21,130 *g* for 30 min at 4°C and washed twice with ice-cold acetone. Final protein pellets were resuspended in 1x SDS-PAGE sample buffer (prepared by diluting the 5x stock in water).

### Metabolic pulse-labelling of nascent proteins

#### Engineered K562 cell lines

Following at least two passages in RPMI^+S^, cells were pelleted and seeded at a density of 200,000 cells/mL in either HPLM^+dS^ or RPMI^+dS^. After 48 h incubation, cells were pelleted and re-seeded to the same initial density in the respective parent medium. After 48 h incubation, cells were pelleted and seeded at a final density of 250,000 cells/mL in 10-cm dishes containing 16 mL of the appropriate medium. After 16-18 h incubation, cells were washed once with 1 mL warm PBS and then seeded in 10-cm dishes containing 16 mL of the corresponding methionine-free medium (HPLM-AHA or RPMI-AHA, see Supplementary Table 1). Following a 1-h incubation, L-Azidohomoalanine (AHA) (50 μM) (Thermo Fisher C10102) was added to each culture and the plates were gently agitated. After an additional 30 min incubation, cells were collected and washed twice with warm PBS and then lysed according to the manufacturer’s protocol (Click-IT Metabolic Labeling Reagents for Proteins, Molecular Probes 10186). Cell lysates were cleared by centrifugation at 21,130 *g* for 10 min at 4°C and then quantified for protein concentration (see **Cell lysis for immunoblotting**). For copper-catalyzed click chemistry, 200 μg of protein was reacted with 40 μM tetramethylrhodamine (TAMRA) alkyne (Thermo Fisher T10183) in light-protected microcentrifuge tubes using the Click-IT Protein Reaction Buffer Kit (Thermo Fisher C10276) for 20 min at room temperature with rotation, according to the manufacturer’s protocol.

Proteins were precipitated according to the manufacturer’s instructions (Click-IT Metabolic Labeling Reagents for Proteins, Molecular Probes 10186). The resulting pellets were resuspended in 200 μL of 1x SDS-PAGE sample buffer (prepared by diluting the 5x stock in PBS). Following 10 min of vortexing and incubation at 70°C for 10 min, samples were centrifuged at 21,130 *g* for 1 min to collect supernatants. Aliquots were loaded onto parallel SDS-PAGE gels for anti-TAMRA immunoblotting and total protein visualization using SimplyBlue SafeStain (Thermo Fisher LC6060). TAMRA signal and total protein abundance were quantified using Image Studio 5.2 (LI-COR).

### ER-phagy flux analysis

Following at least two passages in RPMI^+S^, cells expressing mCherry-eGFP-RAMP4 were pelleted and seeded at a density of 500,000 cells/mL into 6-cm dishes containing 4 mL of the following: (1) EBSS (Thermo Fisher 24010043); (2) complete HPLM^+dS^; (3) alanine-free HPLM^+dS^; (4) RPMI^+dS^; or (5) RPMI^+dS^ supplemented with 430 μM alanine. After 24 h incubation, cells were pelleted and resuspended in 1 mL warm PBS. Samples were then evaluated for eGFP and mCherry signals on an Attune NxT BRV6Y Flow Cytometer (Thermo Fisher). Attune Cytometric Software (Thermo Fisher) was used to collect data. Gating and analysis for ER-phagy flux was performed using FlowJo 10.10 (BD Biosciences) as previously described^61^.

### Quantitative real-time PCR (qPCR)

Following at least two passages in RPMI^+S^, cells were harvested by centrifugation at 250 *g* for 5 min and washed once with 1 mL PBS. Cell pellets were then resuspended in QIAzol lysis reagent (QIAGEN 79306) and homogenized using QIAshredder columns (QIAGEN 79656). RNA was extracted using the miRNeasy Mini Kit (QIAGEN 217004) according to manufacturer’s instructions. For cDNA synthesis, 2.5 μg of RNA was reverse-transcribed using the SuperScript IV First-Strand Synthesis System (Thermo Fisher 18091050) with oligo (dT)20 primers, according to the manufacturer’s protocol. qPCR was performed using PowerUp SYBR Green Master Mix (Thermo Fisher A25742) with the following primer pairs: ACTB_qPCR-F/ACTB_qPCR-R or GPT2_qPCR-F/GPT2_qPCR-R. Reactions were carried out on a CFX Opus 96 Real-Time PCR system (Bio-Rad), and relative expression was performed using CFX Maestro (Bio-Rad, v. 5.3), via the ΔCq method.

### Flag-based affinity purification of recombinant proteins

#### For immunoblot analysis

Following at least two passages in RPMI^+S^, cells expressing Flag-fused cDNAs were pelleted and resuspended to a density of 200,000 cells/mL in either HPLM^+dS^ or RPMI^+dS^. After 48 h incubation, cells were pelleted and seeded at a density of 150,000 cells/mL in 75 mL of the respective parent medium across each of three T-175 flasks. Following another 72-h incubation, 30 million cells were centrifuged at 250 *g* for 5 min, washed once with 1 mL ice-cold PBS, and lysed in 200 μL lysis buffer (see **Cell lysis for immunoblotting**). Cell lysates were cleared by centrifugation at 21,130 *g* for 10 min at 4°C. For anti-FLAG affinity purification (AP), FLAG-M2 affinity gel (Sigma-Aldrich) was washed three times in ice-cold lysis buffer. 100 μL of a 50:50 affinity gel slurry was added to 150 μL of clarified lysate and incubated with rotation for 3 h at 4°C. Following incubation, the beads were washed three times with wash-1 buffer (lysis buffer without protease inhibitor cocktail) and then two times with wash-2 buffer (wash-1 buffer with NaCl adjusted to 500 mM). Recombinant protein was eluted in wash-1 buffer containing 3x-FLAG peptide (200 μg/mL) for 30 min with rotation at room temperature. The eluent was isolated by centrifugation at 100 *g* for 4 min at 4°C using Micro Bio-Spin chromatography columns (Bio-Rad 732-6204) and then denatured by the addition of 5x SDS-PAGE sample buffer.

#### For subsequent mass spectrometry (MS)

The Flag-based AP procedure was identical to that above with minor modifications:

1. Lysis buffer: 20 mM Tris-HCl pH 7.4, 0.1% Digitonin, 100 mM NaCl, 5 mM MgCl2, 1 mM DTT, 1 tablet of PhosStop phosphatase inhibitor cocktail (Roche 04906845001; per 10-mL buffer), and 1 tablet of EDTA-free protease inhibitor (per 25-mL buffer).
2. Wash-1 buffer: Lysis buffer without the protease and phosphatase inhibitor tablets.
3. Wash-2 buffer: Wash-1 buffer without digitonin.
4. Flag eluents were frozen at -80°C until processing for subsequent MS analysis rather than denatured in 5x SDS-PAGE sample buffer.

### Proteomic analysis of Flag-purified recombinant proteins

Flag-purified eluents (1-4 μg protein) were digested with 0.2 μg sequencing-grade trypsin for 16-18 h at 37°C. Reactions were quenched and acidified with 1% trifluoroacetic acid (TFA) prior to desalting. Peptides were desalted using C18-StageTips, constructed by double-layering CDS Empore 2215 C18-HD solid phase extraction disks (CDS Analytical 98-0604-0217-3EA) into P200 pipette tips. StageTips were conditioned with 50 μL acetonitrile and equilibrated with 50 μL of 0.1% TFA in Optima-grade water, with centrifugation at 1,000 *g* for 1 min after each step. Acidified samples were loaded, and tips were washed twice with 50 μL of 0.1% TFA, once with 0.1% TFA in 5% methanol, and once with 0.1% formic acid in 2% acetonitrile. Desalted peptides were eluted in three 20 μL fractions of 0.1% formic acid in 50% acetonitrile. Combined fractions (60 μL total) were concentrated to dryness in a vacuum centrifuge and then stored at -80°C until LC-MS/MS analysis.

#### LC-MS/MS Data Collection

Dried peptide samples were reconstituted in 20 μL of 0.1% formic acid. Total peptide concentration was determined using the Pierce Quantitative Colorimetric Peptide Assay (Thermo Fisher 23275). For each analysis, 300 ng of peptide was loaded onto an Acclaim PepMapRSLC column (75 μm x 50 mm, 2 μm particle size, 100 Å pore size) using an UltiMate 3000 RSLCnano liquid chromatography system coupled to an Orbitrap Exploris 480 hybrid quadrupole-Orbitrap mass spectrometer (Thermo Fisher Scientific). Peptides were loaded over 15 min at 0.3 μL/min in 97% mobile phase A (0.1% formic acid) and 3% mobile phase B (80% acetonitrile, 0.1% formic acid). Peptides were eluted at 0.3 μL/min using a linear gradient from 3% to 50% B over 60 min. Peptides were ionized through a nanospray emitter at 2,000 V using a DirectJunction adapter.

Full MS1 scans were acquired at 60,000 resolution (at 200 m/z) over a range of 350-1,200 m/z with a Normalized AGC Target of 300% (3 x 10^6^ absolute AGC) and the Maximum Injection Time set to ‘Auto’. The top 20 most abundant precursors (charge states 2-6) were selected for MS/MS analysis with an isolation window of 1.4 m/z and a precursor intensity threshold of 5 x 10^3^. Dynamic exclusion was set to 20 s with a precursor mass tolerance of ±10 ppm. MS2 scans were performed using HCD fragmentation (30% normalized collision energy) and acquired at 15,000 resolution (at 200 m/z) with a fixed first mass of 110 m/z. The AGC target was set to ‘Standard’ with a maximum injection time of 22 ms.

### Label-free quantitative proteomics

Following at least two passages in RPMI^+S^, cells were pelleted and resuspended to a density of 200,000 cells/mL in either HPLM^+dS^ or RPMI^+dS^. After 48 h incubation, cells were pelleted and seeded at a density of 250,000 cells/mL in 35 mL of the respective parent medium across each of six T-75 flasks. After 16-18 h incubation, 10 million cells were pelleted, washed twice in ice-cold PBS, snap-frozen in liquid nitrogen and stored at -80°C until processing.

Cell pellets were thawed on ice, reconstituted in 150 µl of 50 mM ammonium bicarbonate, and lysed by passage through a 23-gauge needle. Total protein was quantified using the MicroBCA Protein Assay Kit (Thermo Fisher 23235). A total of 20 μg soluble protein was sequentially incubated with 10 mM DTT for 30 min at 37°C and 55 mM iodoacetamide for 45 min at 37°C in a final volume of 150 µl. Sequencing-grade trypsin (0.5 µg) was added to each sample and incubated at 37°C for 4 h, followed by the addition of another 0.5 µg of trypsin for overnight digestion at 37°C. Reactions were then dried under vacuum (Thermo Savant SPD140DDA) for 1 h at 45°C. Peptides were desalted using ZipTip C18 tips (Millipore ZTC18S) according to the manufacturer’s protocol.

#### LC-MS/MS Data Collection

A total of 2.0 µL of each sample was analyzed using a Waters ACQUITY UPLC M-Class system coupled to a timsTOF flex mass spectrometer (Bruker Scientific) via a CaptiveSpray nano-ESI source. Peptides were trapped on a Waters NanoEase M/Z Symmetry C18 column (180 µm x 20 mm, 5 µm particle size, 100 Å pore size) at a flow rate of 30 μL/min and separated on a Waters NanoEase M/Z Peptide BEH C18 column (75 µm x 150 mm, 1.7 µm particle, 300 Å pore size). Buffer A was 0.1% formic acid in water and Buffer B was 0.1% formic acid in acetonitrile. Peptides were eluted at 0.43 μL/min using a linear gradient from 1% to 95% B over 80 min.

The mass spectrometer was operated in positive ion PASEF mode. The capillary voltage was set to 1500 V. MS and MS/MS spectra were acquired with a scan range of 100-1,700 m/z. The ion mobility (1/K0) was ramped from 0.7 to 1.43 V·s/cm^2^ with a 100 ms accumulation/ramp time and a 100% duty cycle. For PASEF MS/MS, charge states 0-5 were selected for fragmentation. The total cycle time was 1.17 s, comprising one MS1 scan and 10 PASEF MS/MS ramps, with a target intensity of 14,500 and a minimum threshold of 1,750. Collision energy was ramped as a function of mobility, ranging from 20 eV to 59 eV.

### Statistical Analysis

*P* values for all comparisons were determined using a two-tailed Welch’s *t*-test. Statistical analyses were performed with Microsoft Excel and GraphPad Prism. The exact value of *n* and the definition of center (mean) and precision (s.d. or s.e.m.) are described in the figure legends. Bar graphs and plots were prepared in GraphPad Prism. Densitometric quantifications for metabolic pulse-labeling and immunoblot bands were carried out using LI-COR Image Studio (v. 5.2).

#### Proteomic analysis of Flag-purified recombinant proteins

Following data acquisition, Thermo RAW files were searched against the human SwissProt database using the SEQUEST algorithm in Proteome Discoverer v. 2.4 (Thermo Fisher). The precursor mass tolerance was set to 10 ppm and the fragment mass tolerance was set to 0.02 Da. Search parameters included carbamidomethylation at Cys (+57.021 Da) as a static modification and the following dynamic modifications: oxidation at Met (+15.995 Da), acetylation at protein N-termini (+42.011 Da), Met loss at protein N-termini (-131.040 Da), Met loss + acetylation at protein N-termini (-89.030 Da), and GlyVal at Lys (+156.090 Da). Cleavage specificity was set to trypsin with up to two missed cleavages allowed. The Percolator node was used for PSM validation at a false discovery rate (FDR) of 1%. The Proteins results tables in Proteome Discoverer were exported for downstream analysis.

For each condition (HPLM^+dS^ or RPMI^+dS^), Peptide-Spectrum Match (PSM) counts for each prey were normalized to the total PSM counts per sample and then scaled by the mean total PSM count across all nine samples in that condition to generate normalized PSM (nPSM) values. Fold change values were calculated as the ratio of the mean nPSM for each UFM1 variant bait to the mean nPSM of the RAP2A control, following the addition of a 0.5 pseudocount to all means.

To assess the significance of protein-protein interactions for each condition, raw PSM counts were uploaded to the REPRINT^91^ analysis pipeline. RAP2A data were designated as the negative control. Analysis was performed using the SAINT^92^ option with default settings to calculate the False Discovery Rate (FDR). Prey proteins were then filtered to require at least 2 unique peptides across all replicates of the respective UFM1 variant. For enriched proteins, log2 (fold change) was plotted against -log(FDR). For prey with an FDR of 0, the value was capped at y = 5 to facilitate plotting.

#### Label-free quantitative proteomics

Following data acquisition, raw files were searched against the UniProt *H. sapiens* reference proteome using FragPipe v. 23.1 powered by the MS Fragger v. 4.3 search engine. The search used a target-decoy database to estimate the false discovery rate (FDR). Post-processing and validation were performed using Philosopher v. 5.1 and ProteinProphet, with filtering criteria set to achieve a 1% FDR used with the standard settings at the protein level.

Differential abundance analysis was performed using FragPipe-Analyst (v. 1.22)^85^ with the following parameters: standard intensities, variance-stabilizing normalization, Perseus-style imputation, and Benjamini-Hochberg multiple testing correction. The log2 fold change and adjusted *P* values (FDR) for the following comparisons are found in Supplementary Table 4: (1) control cells in HPLM^+dS^ versus RPMI^+dS^; (2) *UFM1*-knockout versus control cells in RPMI^+dS^; and (3) *UFM1*-knockout versus control cells in HPLM^+dS^. To identify significantly enriched biological processes and cellular components among pooled sets of differentially abundant hits, proteins were queried using the DAVID Functional Annotation tool^86,87^. Enrichment was assessed using the Gene Ontology (GO) “Biological Process Direct” and “Cellular Components Direct” annotation datasets, with *H. sapiens* designated as the background proteome. Significance was assessed using the Benjamini-Hochberg adaptive linear step-up method^93^ (reported as FDR in the DAVID pipeline).

#### Focused analysis of label-free quantitative proteomics

For each category, proteins from the final curated lists that were not detected in either FragPipe-Analyst comparison were omitted from the final lists found in Supplementary Table 4.

##### I. Proteins involved in amino acid metabolism

Proteins were initially selected using the REACTOME pathway ‘Metabolism of Amino Acids and Derivatives’ (R-HSA-71291)^94^. This set was refined by manually incorporating three additional proteins (GCAT, PTER, and SHMT2) and excluding the following: cytosolic ribosomal proteins, proteasome subunits, aminoacyl tRNA synthetases, and translation factors.

##### II. Proteins involved in ribosome-associated quality control

Proteins were initially selected using the following REACTOME pathways: ‘Ribosome-associated quality control’ pathway (R-HSA-9948299) and ‘Mitochondrial ribosome-associated quality control’ (R-HSA-9937383)^94^. This set was refined by excluding ribosomal proteins.

##### III. Mitochondrial proteins

Proteins were selected according to inclusion in MitoCarta 3.0^65^.

##### IV. Proteins involved in translation (initiation, elongation, termination)

Proteins were initially selected using the following REACTOME pathways: ‘Eukaryotic translation initiation’ (R-HSA-72613), ‘Eukaryotic translation elongation’ (R-HSA-156842), ‘Eukaryotic translation termination’ (R-HSA-72764), and ‘Mitochondrial translation’ (R-HSA-5368287)^94^. This set was refined by manually incorporating EIF1 and by excluding ribosomal proteins.

##### V. Proteins involved in translation (tRNA aminoacylation)

Proteins were selected using the REACTOME pathway ‘tRNA aminoacylation’ (R-HSA-379724)^94^.

##### VI. Proteins involved in translation (ribosomal proteins)

Proteins were selected using the HGNC groups: ‘S ribosomal proteins’, ‘L ribosomal proteins’, and ‘Mitochondrial ribosomal proteins’^95^.

##### VII. Proteins involved in co-translational protein targeting to ER

Proteins were initially selected using the REACTOME pathway ‘SRP-dependent cotranslational protein targeting to membrane’ (R-HSA-1799339)^94^. This set was refined by excluding ribosomal proteins.

## Data availability

All data needed to evaluate the findings of this study can be found within the article, extended data, or supplementary information. The individual plasmids generated in this study will be deposited in Addgene. AP-MS and quantitative proteomics datasets will be deposited in an appropriate repository. Unique reagents generated in this study will be made available upon reasonable request from the corresponding author.

## ACKNOWLEDGEMENTS

We thank members of the Cantor Lab for upkeep of the LC-MS system and for helpful discussions, A. Mehle and the University of Wisconsin (UW)-Madison Department of Medical Microbiology and Immunology for use of its ultracentrifuge, A. Trinquier and J. Wang for training and access to the BioComp Gradient Station (supported by NIH 3R35GM127088-04S1), M. Pellitteri Hahn for tips on proteomics analysis, and the UW School of Pharmacy Analytical Instrumentation Center for use of its facilities and services (supported by NIH S10-OD028473 to L. Li). This work was supported by grants from the NIH (R35GM156513) and startup funds from the Morgridge Institute for Research (to J.R.C.), and the David and Lucille Packard Fellowship for Science and Engineering (2021-73007) and startup funds from the UW-Madison Department of Biochemistry (to A.M.W.). Fellowship support was provided by the NIH (T32GM152341 to W.E.L.) and the UW-Madison Department of Biochemistry (to G.B.K).

## AUTHOR CONTRIBUTIONS

G.B.K. and J.R.C. initiated the project and designed the research plan. G.B.K. performed most of the experiments, with assistance from S.E.D. W.E.L. performed the Flag affinity purification proteomics experiments with supervision from A.M.W. J.R.C. performed metabolomics experiments. G.B.K. and J.R.C. analyzed and interpreted the experimental data. J.R.C. wrote the manuscript with assistance from G.B.K. All authors discussed the manuscript. J.R.C. supervised the studies.

## COMPETING INTERESTS

J.R.C. is an inventor on an issued patent for Human Plasma-Like Medium assigned to the Whitehead Institute (Patent number: US11453858). The remaining authors declare no competing interests.

Supplementary information is available for this paper (Supplementary Tables 1-5).

---

**Extended Data Fig. 1.**
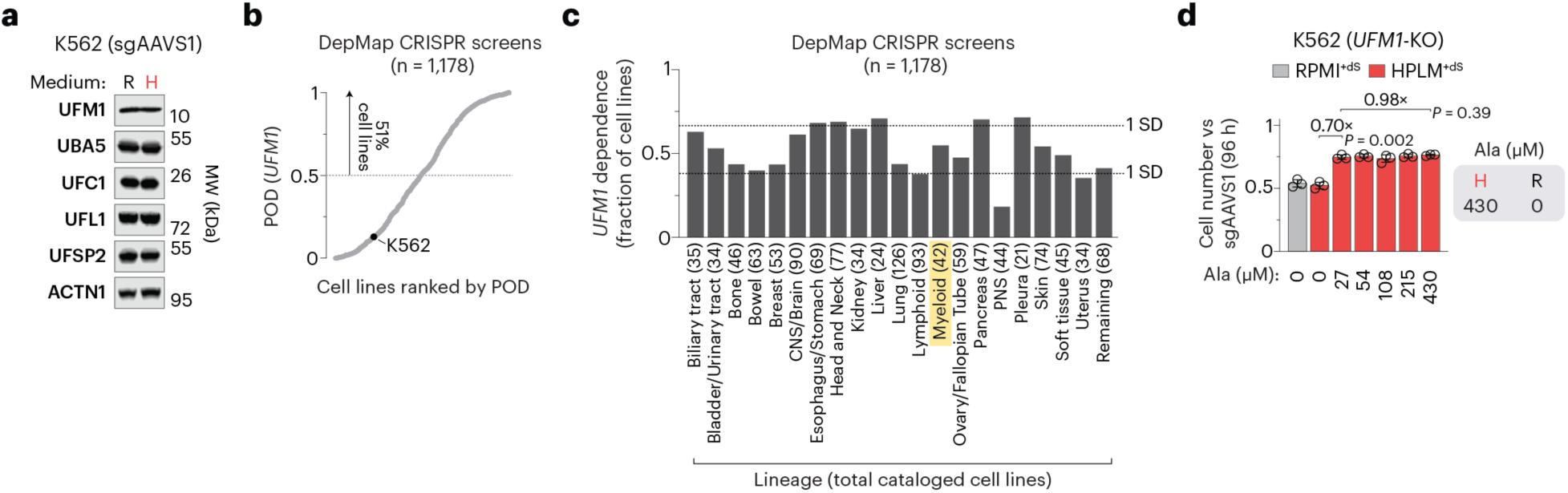
Additional data related to conditional dependence on the UFMylation system is linked to alanine availability. (a) Immunoblots for expression of UFM1, UBA5, UFC1, UFL1, and UFSP2 in control cells. ACTN1 served as the loading control. (b) Human cancer cell lines ranked by probability of dependency for *UFM1* across CRISPR screens in DepMap^7^. Probability > 0.5 is the reference threshold for essentiality. (c) Distribution of *UFM1* dependency across lineages in DepMap. The total of number of cell lines in each lineage is indicated in parentheses. K562 belongs to the myeloid lineage (highlighted). CNS, central nervous system. PNS, peripheral nervous system. (d) Relative growth of *UFM1*-knockout versus control cells (mean ± s.d., *n* = 3 biologically independent samples). Two-tailed Welch’s *t*-test comparing the respective mean ± s.d. (bar) versus mean ± s.d. (control cells) between bars. Values above brackets indicate fold change between bars (left). Defined alanine levels in HPLM and RPMI (right).

**Extended Data Fig. 2.**
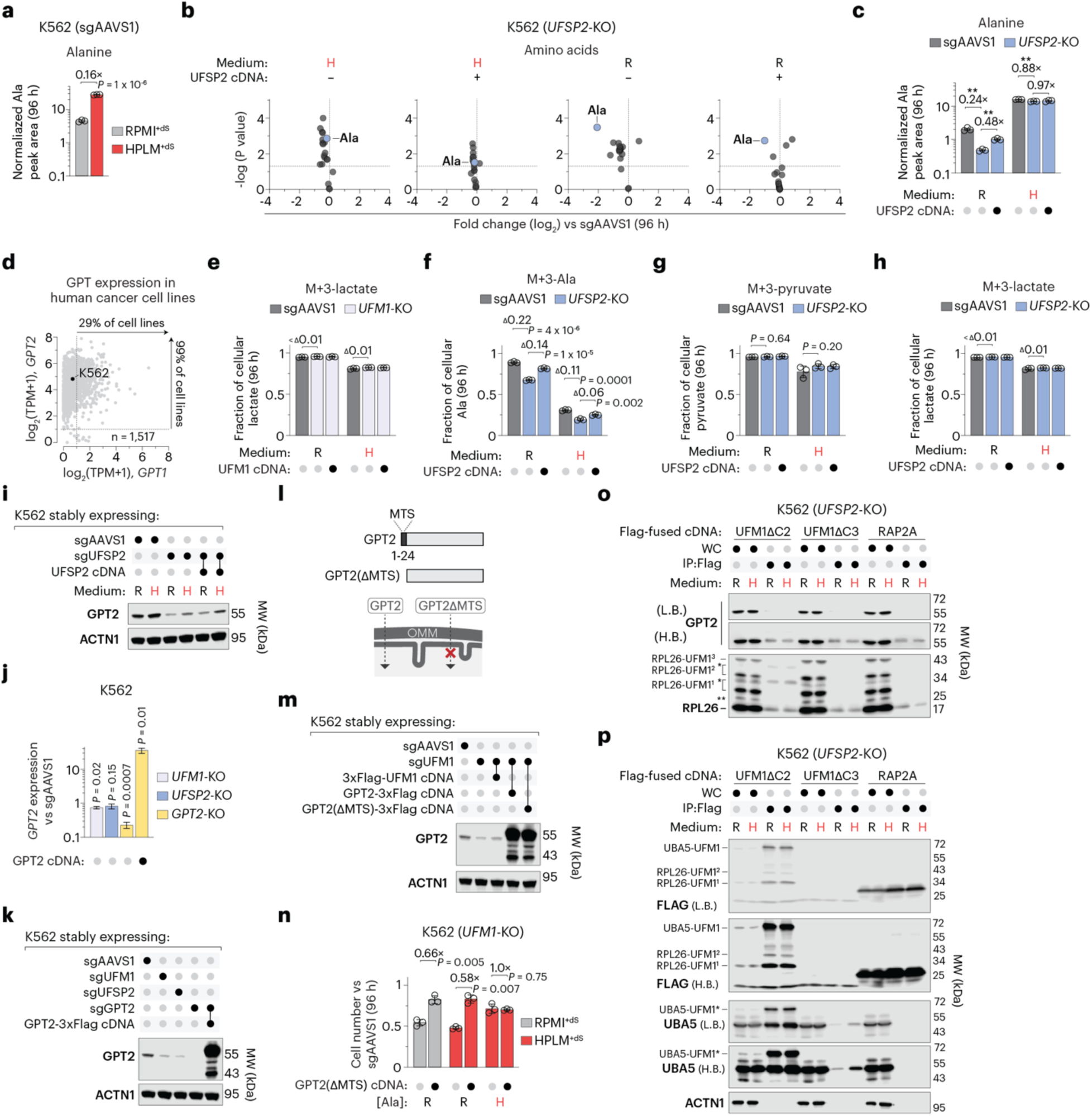
Additional data related to the UFMylation system maintains GPT2 abundance. (a) Alanine levels in control cells (mean ± s.d., *n* = 3 biologically independent samples). Two-tailed Welch’s *t*-test. (b) Relative amino acid levels in *UFSP2*-knockout versus control cells (*n* = 3 biologically independent samples). Horizontal dotted lines mark a *P* value of 0.05 (y-axis). (c) Alanine levels in *UFSP2*-knockout and control cells (mean ± s.d., *n* = 3 biologically independent samples). Two-tailed Welch’s *t*-test. \*\**P* < 0.005. (d) Comparison between *GPT1* and *GPT2* mRNA transcript levels^26^. TPM, transcripts per million. (e) Fractional labeling of lactate in *UFM1*-knockout and control cells (mean ± s.d., *n* = 3 biologically independent samples). Two-tailed Welch’s *t*-test. (f-h) Fractional labeling of alanine (F), pyruvate (G), and lactate (H) in *UFSP2*-knockout and control cells (mean ± s.d., *n* = 3 biologically independent samples). Two-tailed Welch’s *t*-test. (i) Immunoblot for expression of GPT2 in *UFSP2*-knockout and control cells. ACTN1 served as the loading control. (j) Relative *GPT2* mRNA expression quantified by qPCR in the indicated knockout lines versus control cells (mean ± s.d., *n* = 3 biologically independent samples). Two-tailed Welch’s *t*-test. (k, m) Immunoblots for expression of GPT2 in the indicated knockout and control cells. ACTN1 served as the loading control in each case. (l) Schematic of domain architecture for wild-type (top) and MTS-deficient GPT2 (bottom). Arrows indicate predicted mitochondrial translocation. MTS, mitochondrial targeting signal. (n) Relative growth of *UFM1*-knockout versus control cells (mean ± s.d., *n* = 3 biologically independent samples). Two-tailed Welch’s *t*-test comparing the respective mean ± s.d. (bar) versus mean ± s.d. (control cells) between bars. (o, p) Immunoblots of GPT2, RPL26 (o), Flag and UBA5 (p) in whole-cell lysates (WC) and Flag affinity-purified eluents (IP:Flag) from *UFSP2*-knockout cells. ACTN1 served as a loading control for WC. Single asterisks denote Flag-shifted UFMylated species; double asterisks in (o) indicate non-specific bands. L.B., low brightness. H.B., high brightness. In a, c, and n, values above brackets indicate fold change between bars. In e-h, values above brackets indicate differences in fractional labeling between bars.

**Extended Data Fig. 3.**
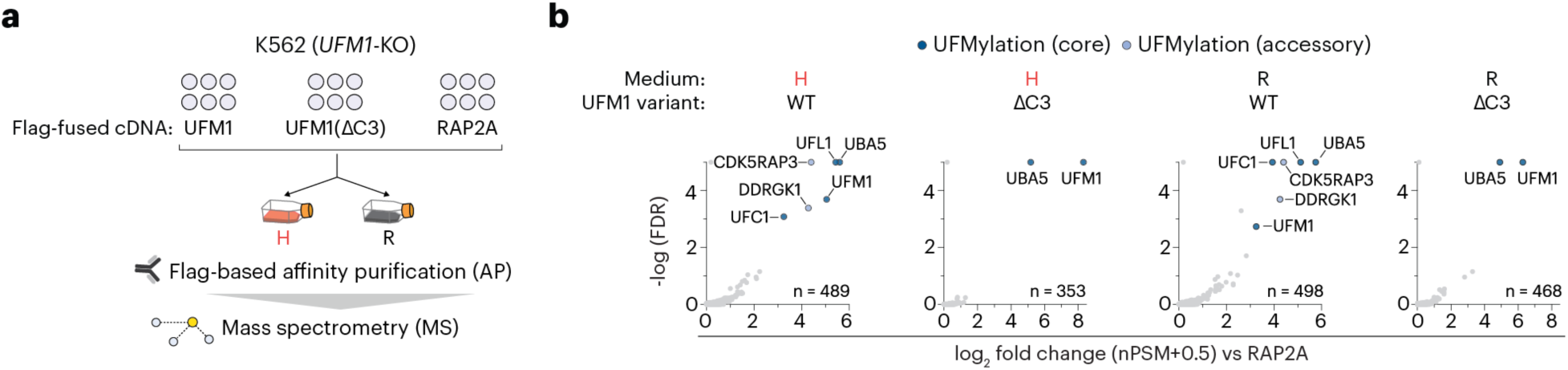
Additional data related to the UFMylation system acts as a ribosome collision counter at the ER. (a) Schematic of the Flag-based affinity purification-mass spectrometry (AP-MS) workflow in *UFM1*-knockout cells. (b) Relative enrichment of prey proteins captured by the indicated 3xFlag-UFM1 variants compared to control bait (RAP2A-3xFlag) (*n* = 3 biologically independent samples). FDR values were determined using SAINT analysis^91,92^. Proteins were subsequently filtered for detection with > 1 unique peptide count across all respective UFM1 variant replicates in the associated condition. nPSM, normalized Peptide-Spectrum Match.

**Extended Data Fig. 4.**
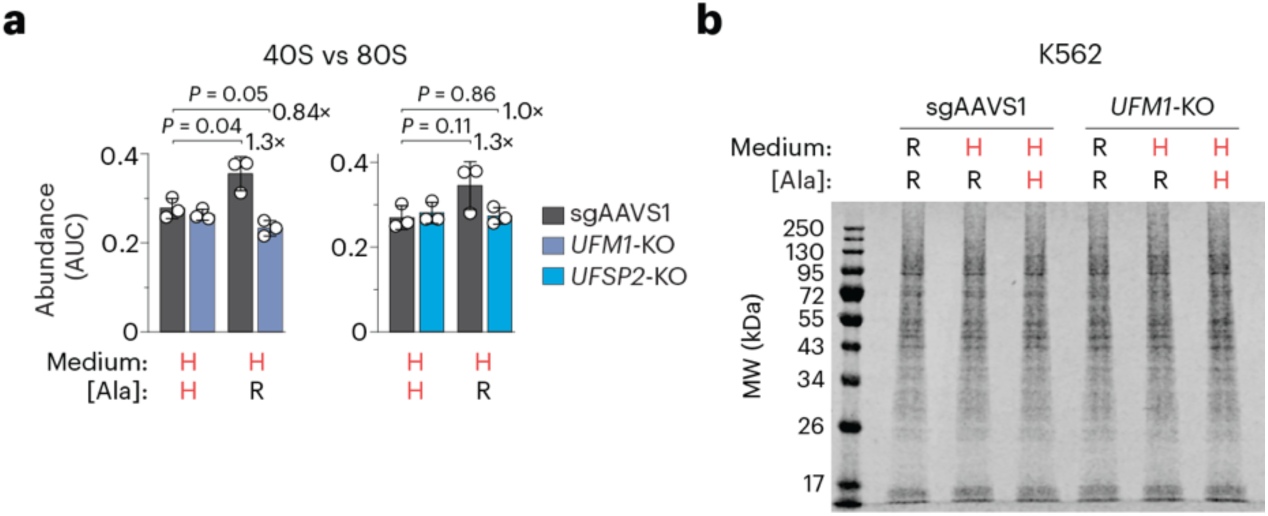
Additional data related to dynamic UFMylation is conditionally required to maintain protein synthesis rates. (a) Ratio of free 40S subunits to 80S monosomes in *UFM1*-knockout, *UFSP2*-knockout, and control cells (mean ± s.d., *n* = 3 biologically independent samples). Two-tailed Welch’s *t*-test. Values beside brackets indicate fold change between bars. (b) Pseudocolor Coomassie-stained gel for total protein levels in lysates from *UFM1*-knockout and control cells, used for total protein normalization of the TAMRA signal shown in Fig. 4j. In a and b, H, HPLM-defined concentration; R, RPMI-defined concentration.

**Extended Data Fig. 5.**
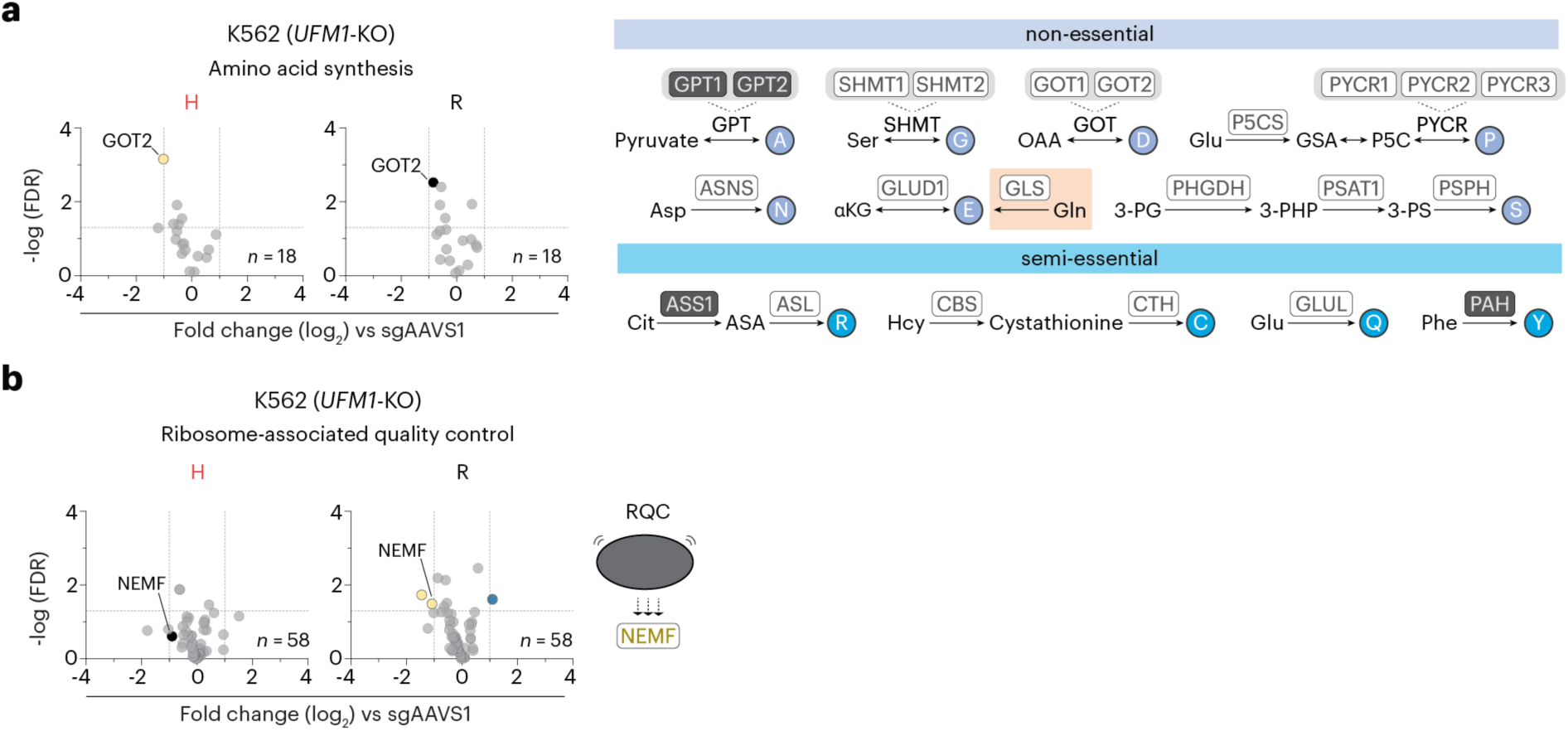
Additional data related to *UFM1* deletion remodels the global proteome. (a, b) Relative protein levels in *UFM1*-knockout versus control cells (*n* = 6 biologically independent samples). Schematic depicts de novo synthesis pathways for non-essential and semi-essential amino acids (a). While not strictly defined as de novo synthesis, the GLS-mediated reaction is included as a major source of glutamate. Dark gray boxes denote enzymes not detected in the proteomics datasets. NEMF responds to ribosome stalling (b). log2 fold change and FDR values were determined using FragPipe-Analyst^85^. Dotted lines denote a fold change of ± 2 (x-axis) and an FDR of 0.05 (y-axis).

**Extended Data Fig. 6.**
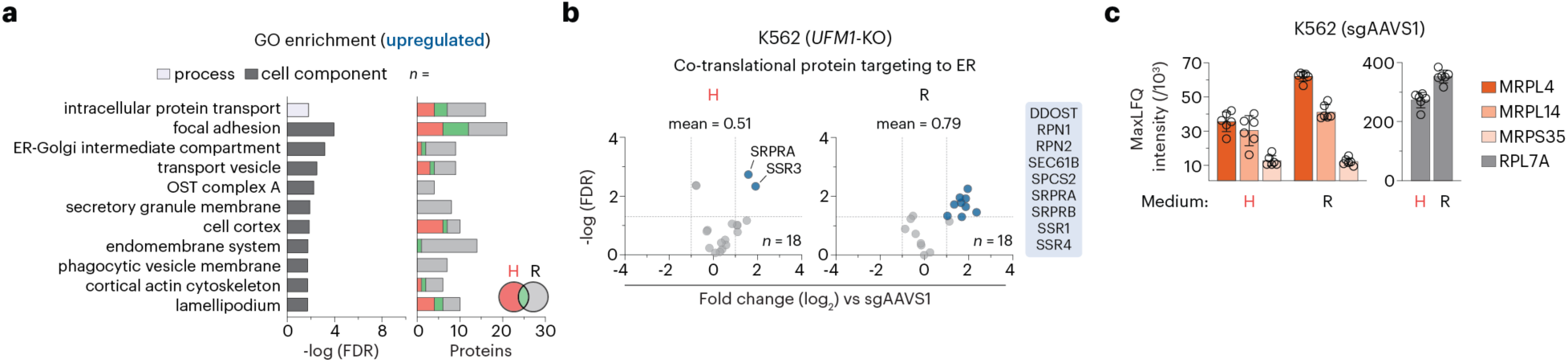
Additional data related to *UFM1* deletion alters mitochondrial proteostasis. (a) Enriched Gene Ontology (GO) biological processes and cellular components identified among the 290 total upregulated protein hits in *UFM1*-knockout versus control cells (left). Categories were identified using DAVID functional annotation analysis^86,87^. Number of proteins per condition in each respective category (right). (b) Relative protein levels in *UFM1*-knockout versus control cells (*n* = 6 biologically independent samples). Mean values represent the average log2 fold change across the displayed protein sets. log2 fold change and FDR values were determined using FragPipe-Analyst^85^. Dotted lines denote a fold change of ± 2 (x-axis) and an FDR of 0.05 (y-axis). (c) Protein abundance in control cells (mean ± s.d., *n* = 6 biologically independent samples). MaxLFQ (Label-Free Quantification) intensities were quantified using FragPipe-Analyst. RPL7A is included for comparison with the three mitoribosomal proteins.

**Extended Data Fig. 7.**
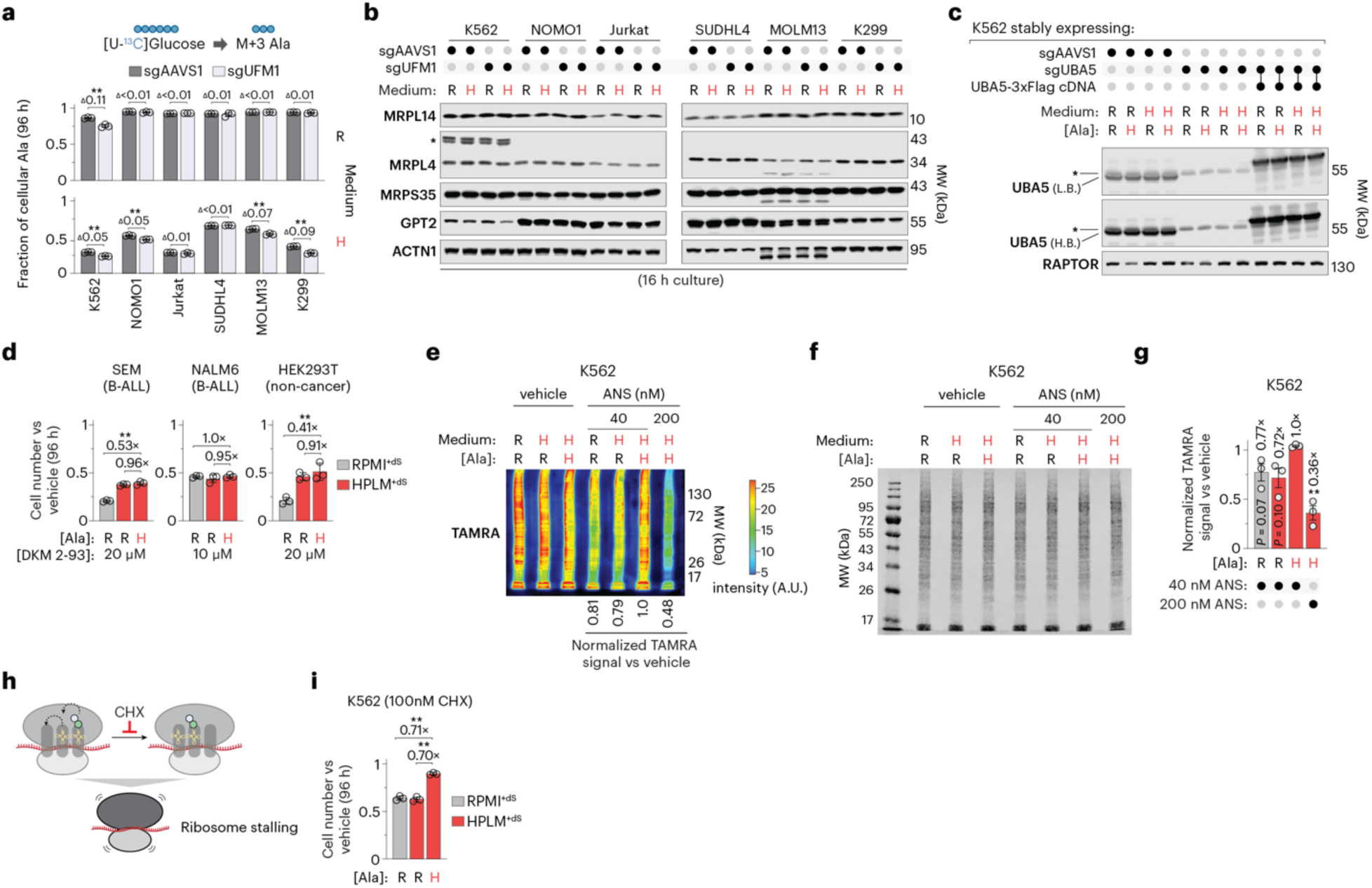
Additional data related to conditional dependence on the UFMylation system varies with cell-intrinsic factors. (a) Fractional labeling of alanine for cell lines transduced with sgUFM1 or sgAAVS1 (mean ± s.d., *n* = 3 biologically independent samples). Two-tailed Welch’s *t*-test. \*\**P* < 0.005. Values above brackets indicate differences in fractional labeling between bars. (b) Immunoblots for expression of MRPL14, MRPL4, MRPS35, and GPT2 in cell lines transduced with either sgAAVS1 or sgUFM1. Lysates were harvested 16 h post-seeding to match the culture duration used for the quantitative proteomics. ACTN1 served as the loading control. Asterisk denotes a band corresponding to a putative monoubiquitinated MRPL4 species (∼10 kDa shift). (c) Immunoblot for expression of UBA5 in *UBA5*-knockout and control cells. RAPTOR served as the loading control. Asterisk denotes a non-specific band. L.B., low brightness; H.B., high brightness. (d) Relative growth of indicated cell lines treated with DKM 2-93 versus vehicle (mean ± s.d., *n* = 3 biologically independent samples). (e) Pseudocolor immunoblot of TAMRA signal in anisomycin (ANS)- and vehicle-treated cells. Signal intensity is represented by the corresponding colorimetric scale. (f) Pseudocolor Coomassie-stained gel of lysates from ANS- and vehicle-treated cells, used for total protein normalization of the TAMRA signal shown in (e). (g) Relative TAMRA signal in ANS- versus vehicle-treated cells (mean ± s.e.m., *n* = 3 biologically independent samples). For each replicate, raw intensities were normalized by total protein and then scaled to the maximum normalized value within that replicate. Two-tailed Welch’s *t*-test. (h) Schematic of cycloheximide (CHX) function. CHX is a global elongation inhibitor that binds to the 60S subunit (E-site). (i) Relative growth of CHX- versus vehicle-treated cells (mean ± s.d., *n* = 3 biologically independent samples). Values above brackets indicate fold change between bars. In d and i, Two-tailed Welch’s *t*-test comparing the respective mean ± s.d. (bar) versus mean ± s.d. (control cells) between bars. \*\**P* < 0.01. In c-g and i, H, HPLM-defined concentration; R, RPMI-defined concentration.

